# Smaug membraneless organelles regulate mitochondrial function

**DOI:** 10.1101/2020.07.31.223099

**Authors:** Ana J. Fernández-Alvarez, María Gabriela Thomas, Malena L. Pascual, Martín Habif, Jerónimo Pimentel, Agustín A. Corbat, João P. Pessoa, Pablo E. La Spina, Lara Boscaglia, Anne Plessis, Maria Carmo-Fonseca, Hernán E. Grecco, Marta Casado, Graciela L. Boccaccio

## Abstract

Smaug is a conserved translational repressor that recognizes specific RNA motifs in a large number of mRNAs, including nuclear transcripts that encode mitochondrial enzymes. Smaug orthologs have been shown to form membraneless organelles (MLOs) in several organisms and cell types. Using single-molecule FISH we show here that SDHB and UQCRC1 mRNAs associate with Smaug1 MLOs in the human cell line U2OS. Simultaneous loss of function of Smaug1 and Smaug2 affects both mitochondrial respiration and mitochondrial network morphology. Deletion of specific Smaug1 protein regions resulted in impaired MLO formation that correlates with mitochondrial defects. In addition, rotenone but not the respiratory chain uncoupling agent CCCP rapidly induces Smaug1 MLO dissolution. Finally, metformin elicits a similar effect on Smaug1 MLOs and provokes the release of bounded mRNAs. We propose that mitochondrial activity affects Smaug1 MLO dynamics, thus allowing for regulation of nuclear mRNAs that encode key mitochondrial proteins.

## Introduction

Smaug orthologs bind transcripts that contain a variety of stem-loops termed Smaug recognition elements (SREs). Several unbiased screens in *Drosophila* allowed the identification of thousands of target mRNAs that are involved in widely diverse cellular processes (Amadei et al., 2015; Aviv et al., 2006; Baez & Boccaccio, 2005; Cao et al., 2020; Chartier et al., 2015; Chen, Dumelie, et al., 2014; Eichhorn et al., 2016; Johnson & Donaldson, 2006; Laver et al., 2012; Nelson et al., 2004; Niu et al., 2017; Oberstrass et al., 2006; Pinder & Smibert, 2013; Rouget et al., 2010; Semotok et al., 2005; Semotok et al., 2008; She et al., 2017; Tadros et al., 2007). Messenger RNA regulation by Smaug can involve translational repression and/or decay. Both insects and vertebrates Smaug forms membraneless organelles (MLOs) termed Smaug-foci or Smaug-bodies, which contain repressed mRNAs. Smaug bodies were identified in mammalian neuroblasts, hippocampal dendrites, fly embryo and adult muscle, and are distinct from Processing Bodies (PBs), a well-known type of MLO that also contain repressed mRNAs (Amadei et al., 2015; Baez & Boccaccio, 2005; Baez et al., 2011; Chartier et al., 2015; Luchelli et al., 2015; Rouget et al., 2010; Sachdev et al., 2019). The molecular consequences of Smaug-body formation are incipiently described. Condensation of the yeast ortholog Vts1 facilitates the degradation of target mRNAs (Chakravarty et al., 2020), whereas mammalian Smaug1-body dissolution is linked to mRNA release and translational activation of dendritic mRNAs (Baez et al., 2011; Luchelli et al., 2015). The formation of MLOs and related molecular condensates is thought to involve liquid-liquid phase separation (LLPS) processes driven by multiple protein-protein interactions, which remain unknown in the case of Smaug (Courchaine et al., 2016; Guo & Shorter, 2015; Perez-Pepe et al., 2018; Sachdev et al., 2019; Van Treeck et al., 2018).

The dysregulation of Smaug orthologs in different organisms and tissues leads to a wide diversity of phenotypes. Drosophila Smaug loss of function provokes embryo defects due to the abnormal expression of maternal mRNAs. In addition, fly Smaug binds numerous mRNAs that code glycolytic and mitochondrial enzymes (Tadros et al., 2007). The two mammalian paralogs termed Smaug1/Samd4A and Smaug2/Samd4B are involved in neuron differentiation, synapse plasticity and bone development (Amadei et al., 2015; Baez et al., 2011; Luchelli et al., 2015; Niu et al., 2017). Overexpression of *Drosophila* Smaug or of mammalian Smaug1 in muscular cells suppress myotonic dystrophy-1 (MD1) (de Haro et al., 2013), whereas independent work identified both *Drosophila* Smaug and mammalian Smaug1 as enhancers of a mitochondrial defect caused by aberrant mRNA polyadenylation and linked to oculopharyngeal muscular dystrophy (OPMD) (Chartier et al., 2015). Finally, a point mutation in murine Smaug1 generates a complex phenotype characterized by leanness, resistance to fat intake-induced obesity, reduced muscle and adipose tissue, and abnormal morphology of both myofibers and adipocytes (Chen, Holland, et al., 2014). Work in yeast showed that Vts1 is involved in nutrient sensing (Chakravarty et al., 2020; She et al., 2017) and thus, a connection between Smaug and the energetic metabolism appears conserved from yeast to animals.

Here we show that mammalian Smaug1 MLOs interact with nuclear mRNAs that encode mitochondrial enzymes. Specifically, single molecule FISH revealed that succinate dehydrogenase subunit B (SDHB) and ubiquinol-cytochrome C reductase core protein 1 (UQCRC1) mRNAs associated with Smaug1-bodies. In addition, loss of function of Smaug1 and Smaug2 affected both mitochondrial respiration and mitochondrial network complexity. The phenotype was rescued by full-length Smaug1 but not by Smaug1 deletion mutants with defective MLO formation. Finally, exposure to metformin –a metabolism activator widely used for the treatment of type II diabetes– immediately induced Smaug1-MLO dissolution and mRNA release. Complex I inhibition by rotenone similarly induced Smaug1-body dissolution whereas strong uncoupling by exposure to CCCP elicited no effect. We speculate that the condensation of Smaug1 MLOs help the control of nuclear transcripts that encode mitochondrial proteins through the regulated release of mRNAs.

## Results

### SDHB and UQCRC1 mRNAs associate to Smaug1 membraneless organelles

Smaug orthologs are expressed in several mammalian cell lines (Fernandez-Alvarez et al., 2016) and as expected, we found that Smaug1 forms cytosolic puncta in U2OS cells (Figure 1A). Smaug1-bodies were present is most cells (70-80%, 5 independent experiments, N≥200 cells each) with highly variable numbers. Their size and distribution remind that of PBs, however Smaug1-bodies did not colocalize with PBs identified by DCP1A staining (Figure 1A) as previously reported in neurons (Baez et al., 2011). Finally, Smaug1-bodies dissolve upon exposure to cycloheximide, suggesting that they can release translation-competent mRNAs (Figure 1B).

**Figure 1.**
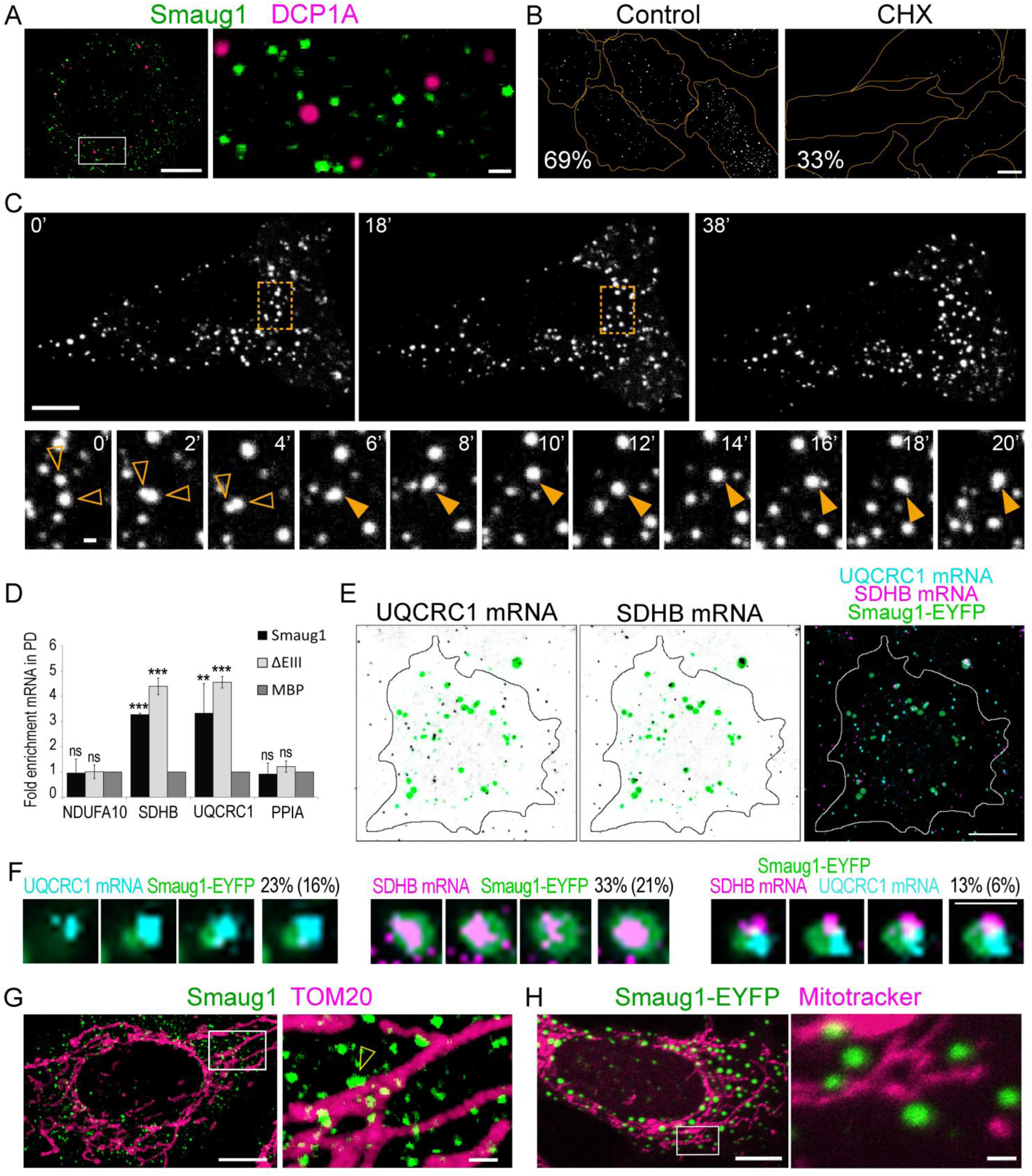
Smaug1-bodies contain mRNAs that encode mitochondrial enzymes. **(A)** U2OS cells were immunostained for DCP1A and Smaug1. **(B)** Cells were exposed to 1µM cycloheximide for one hour. Representative cells from each condition are depicted. The proportion of cells with Smaug1-bodies evaluated as described in Materials and Methods is indicated. Cell contours (orange) were drawn as described in Materials and Methods. **(C)** Smaug1-EYFP was recorded in live cells. Inset, two bodies (empty arrowheads) start merging at 2 min and remain fussed in a single body (full arrowhead). Fusion with a third body starts at 18 min. **(D)** RNA pull-down. Smaug1, Smaug1 ΔEIII or MBP as a negative control, all of them tagged with V5-SBP were pulled-down and the presence of the indicated transcripts in the pulled-down material was analyzed by qRT-PCR. Mean values from three independent experiments of pull-down/input ratio for each transcript normalized to ribosomal protein RpLp0 are plotted. Error bars, standard deviation. Expression levels and recovery of V5-SBP-tagged constructs were confirmed to be similar by western blot (not shown). **(E)** Single molecule FISH for SDHB and UQCRC1 mRNAs was performed in cells expressing Smaug1 EYFP. Cell contour (solid line) was drawn as described in Materials and Methods. **(F)** Three adjacent confocal slices and maximal projection of representative Smaug1-EYFP bodies containing SDHB mRNA, UQCRC1 mRNA, or both transcripts are depicted. Colocalization was analyzed in 400 Smaug1-ECFP bodies from randomly-selected cells and indicated as percentage. Stochastic colocalization (in brackets) was assessed in randomized images as described in Materials and Methods. **(G)** Smaug1 and TOM20 were immunostained. Arrowheads, examples of Smaug1-bodies that contact mitochondria. **(H)** Smaug1-EYFP transfected U2OS cells were live-stained with MitoTracker™ Red CMXRos. Scale bars: whole cells, 10 μm, insets, 1 μm.

Transfected Smaug1-EYFP or Smaug1-V5 condense in cytosolic bodies (Baez & Boccaccio, 2005; Baez et al., 2011) and here we confirmed that they were similar in size and distribution to endogenous bodies. Smaug1-EYFP bodies exclude ribosomal subunits detected by FISH with specific probes for 18S rRNA, indicating that they are not stress granules (Supplementary file 1). In addition, Smaug1-bodies did not contain ubiquitin, a marker of protein aggregates (Supplementary file 1), and thus we conclude that transfection of Smaug1 is a reliable strategy for further studies. Next, we performed real time microscopy and found that Smaug1-EYFP bodies were highly dynamic and they sporadically fuse, a feature characteristic of MLOs (Figure 1C, Video1).

Drosophila Smaug was shown to bind several nuclear transcripts that encode mitochondrial enzymes (Bruzzone et al., 2020; Chartier et al., 2015; Tadros et al., 2007) and among many other putative mammalian targets, we analyzed NADH:ubiquinone oxidoreductase subunit A10 (NDUFA10), succinate dehydrogenase subunit B (SDHB), and ubiquinol-cytochrome C reductase core protein 1 (UQCRC1) mRNAs. We transfected and pulled-down Smaug1 fused to a V5-SBP double tag and evaluated the co-purification of these transcripts by RT-PCR (Figure 1D). Both SDHB and UQCRC1 mRNAs were recovered in the V5-SBP-Smaug1 pull-down material whereas the control construct V5-SBP-MBP did not pull-down these transcripts. In contrast, we found that NDUFA10 mRNA was not significantly enriched and recovery values were close to those of PP1A mRNA (Figure 1D). Finally, Smaug1 ΔEIII, a natural splicing variant that forms similar cytosolic bodies (Fernandez-Alvarez et al., 2016) showed a similar profile. These results are comparable to observations reported in Drosophila, where SDHB mRNA is four-fold enriched in fly Smaug pull-downs and NDUFA10 mRNA is among the less-bounded transcripts (Chartier et al., 2015).

Then, we used single-molecule FISH to simultaneously detect SDHB mRNA, UQCRC1 mRNA and Smaug1-EYFP bodies (Figure 1E). We found that a fraction of Smaug1-bodies contains or contact SDHB or UQCRC1 mRNA puncta. Random colocalization was estimated as described in Material and Methods and resulted lower in both cases (Figure 1F). The presence of SDHB and UQCRC1 mRNAs in a single Smaug1-EYFP-body was not mutually exclusive and on the contrary, we found that the co-occurrence of the two transcripts was slightly higher than that predicted as independent events (0.13 vs 0.33 × 0.23=0.08 in Figure 1F). In addition, we found that Smaug1-bodies are frequently in contact with mitochondria identified by TOM20 immunostaining or by MitoTracker™ Red CMXRos staining (Figure 1G-H). Frequency of contacts with the mitochondria was moderately higher than that calculated in randomized images obtained as previously described for purinosomes (58% SD=9, vs 43% SD=5, N=10) (French et al., 2016).

### Smaug1 and Smaug2 knockdown impairs mitochondrial respiration

We have previously shown that transfected Smaug1 and Smaug2 form cytosolic bodies that colocalize (Fernandez-Alvarez et al., 2016) and we speculated that mammalian Smaug proteins may affect mitochondria activity. We investigated the effect of the double knockdown (KD) of Smaug1 and Smaug2, which were efficiently silenced by specific siRNAs (Figure 2A).

**Figure 2.**
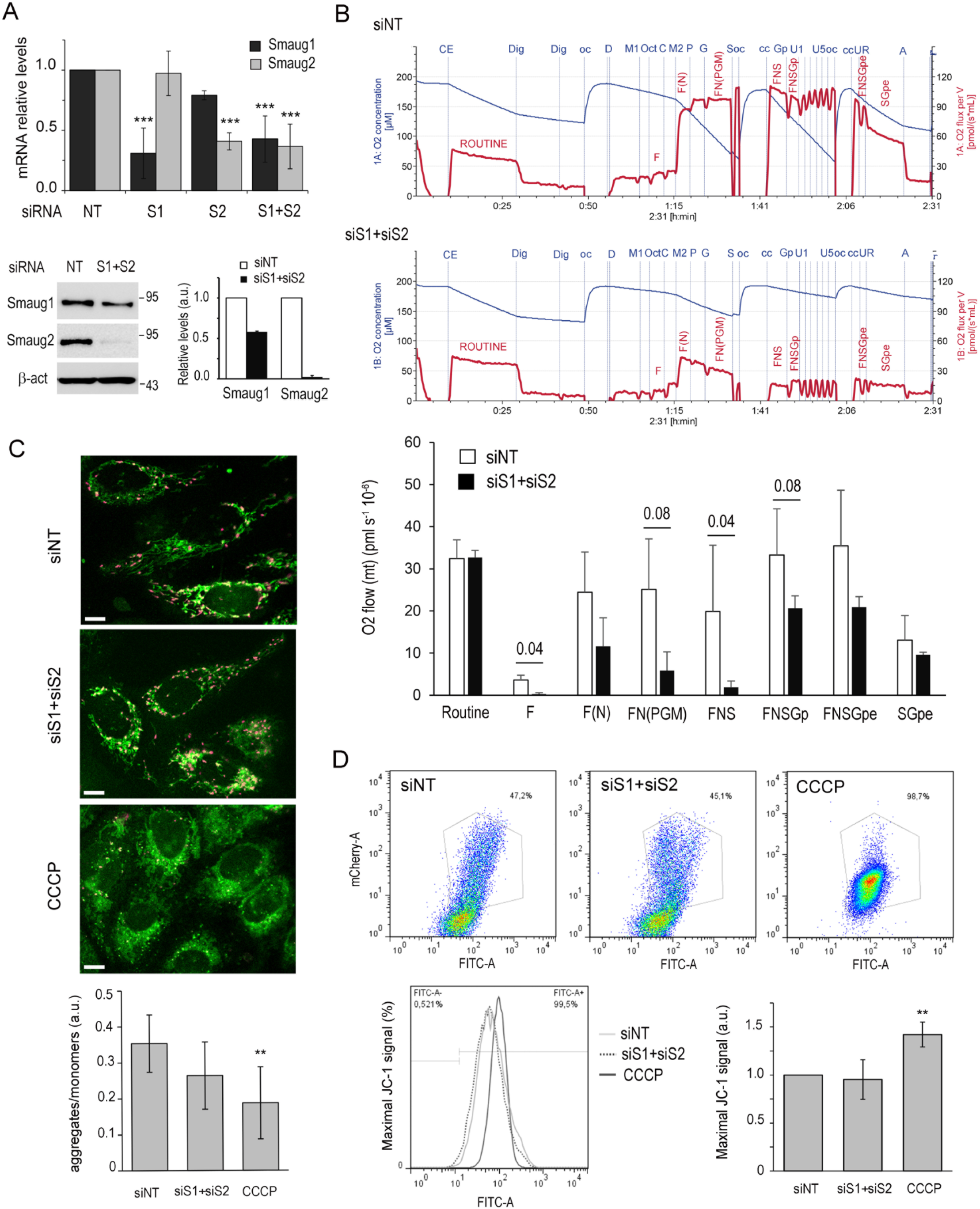
Respiratory defects upon Smaug1 and Smaug2 knockdown. **(A)** U2OS cells were treated with siRNAs against Smaug1 (S1) and/or Smaug2 (S2) during 48 hs and mRNA and protein levels were respectively analyzed by qRT-PCR and western blot. A representative qRT-PCR out of three is shown and two independent experiments were averaged for quantification by western blot. NT, non-targeting siRNA. **(B)** High resolution respirometry in permeabilized cells. Top, representative traces of oxygen concentration (blue) and oxygen flow (red). The addition of digitonin (Dig), ADP (D), 0.1 mM malate (M1), octoanylcarnitine (Oct), cytochrome c (C), 2 mM malate (M2), pyruvate (P), glutamate (G), succinate (S), phosphoglycerate (Gp), CCCP (U1-U5), rotenone (R) and antimycin A (A) is indicated with dotted lines as well as re-oxygenation and closing of the chambers (oc and cc, respectively). Sections reflecting OXPHOS fingerprints (see Methods) are labelled. Bottom, oxygen flow was measured as described in Material and Methods. Mean oxygen flow per million cells ± SE from three independent experiments performed in duplicate are shown. Statistical significance was calculated as described in Materials and Methods, p values lower than 0.1 are indicated. **(C-D)** Mitochondrial membrane potential was determined using JC-1 dye. **(C)** J-aggregates (shown in magenta) and monomers (green) were analyzed by confocal microscopy. The proportion of aggregates/monomers calculated as the average intensity ratio from 10 fields from a representative experiment is plotted. **(D)** J-aggregates (red) or monomers (green) were analyzed in the mCherry and FITC channels respectively by flow cytometry. Histogram, overall green fluorescence (FITC). Bottom right, JC-1 monomer signal was quantified using geometric median from three independent experiments and plotted as fold change ± S.E.M. Statistical analysis was done using Student’s t-test (paired, one tailed) (*P<0.05).

We measured the respiratory capacity of intact or permeabilized cells using high-resolution respirometry as described in Materials and Methods. We found significant changes in permeabilized cells analyzed by the substrate-uncoupler-inhibitor-titration (SUIT)-RP2 protocol (Doerrier et al., 2018) (Figure 2B) while no effects were observed in intact cells (Supplementary file 2). SUIT-RP2 allows to measure the F-pathway, the electron transport capacity, and the maximum oxidative phosphorylation (OXPHOS) by sequential addition of a number of substrates (see Methods). We found that routine endogenous respiration –which depends on intracellular substrates– remained unchanged upon Smaug1+2 double KD, in accordance with the lack of effect observed in intact cells. In contrast, oxygen flux through fatty acid oxidation was reduced upon Smaug1 and Smaug2 double KD (Figure 2B). A moderate reduction in oxygen flow was observed after activation of the N-pathway with malate (F(N)); the effect was more significant in the presence of malate, pyruvate and glutamate (FN(PGM)), and stronger upon the addition of succinate (FNS), suggesting defective complex I and II activities. Oxygen flow increased significantly in both control and KD cells after addition of glycerophosphate, indicating that glycerophosphate dehydrogenase shuttle (FNSGp) is functional in both cases, with almost normal maximum respiratory capacity and oxygen flow after inhibition of complex I (FNSGpe; SGpe). Finally, complex IV activity was not affected by Smaug1+2 double KD (siNT, 168 and siSmaug1+2, 164 pmol/s*mill cells*mtDNA, see methods).

Altogether, these observations indicate decreased succinate-activated respiration upon Smaug1 and Smaug2 double KD. In addition, we studied the effect of Smaug1+2 KD on mitochondrial membrane potential by JC-1 staining followed by flow cytometry or confocal analysis. Cells treated with the strong depolarized agent carbonyl cyanide 3-chlorophenylhydrazone (CCCP) were analyzed in parallel. We found that depletion of Smaug1+2 did not significantly affect mitochondrial membrane potential (Figure 2C, D).

### Mitochondrial network disruption upon Smaug1 and Smaug2 knockdown

Collectively, the above observations indicate that Smaug1+2 KD affects OXPHOS activity without provoking serious mitochondrial damage. We were interested in investigating the relevance of Smaug MLOs condensation to mitochondrial physiology. MLOs are typically assessed by microscopy and thus, we seek for mitochondrial phenotypes compatible with single-cell imaging. The mitochondrial network is dynamically regulated in connection with mitochondrial respiration changes and we analyzed its overall distribution by TOM20 staining (Figure 3). We classified the cells in two groups as short or elongated branched mitochondria and found that Smaug1 or Smaug2 KD significantly reduced the complexity of the mitochondrial network (Figure 3A). In five independent experiments, we found that 50 to 80 % of the control cells showed elongated mitochondria, and this number was reduced to 20-40% upon Smaug1 and Smaug2 double KD. Single KD of either Smaug1 or Smaug2 elicited a statistically significant effect yet not as strong as double KD (Figure 3B). Mitochondria were also analyzed by staining live cells with MitoTracker™ Red CMXRos and similar results were obtained (Supplementary file 3). The mitochondrial DNA content relative to the nuclear DNA content evaluated by qPCR of the mitochondrial gene for cytochrome C oxidase 2 and of the nuclear gene for succinate dehydrogenase subunit A yielded similar values in all conditions (siNT 1±0.3; siSmaug1, 1.1±0.3; siSmaug2 0.9±0.1; siSmaug1+siSmaug2 1.1±0.3), indicating that mitochondrial mass was not affected.

**Figure 3.**
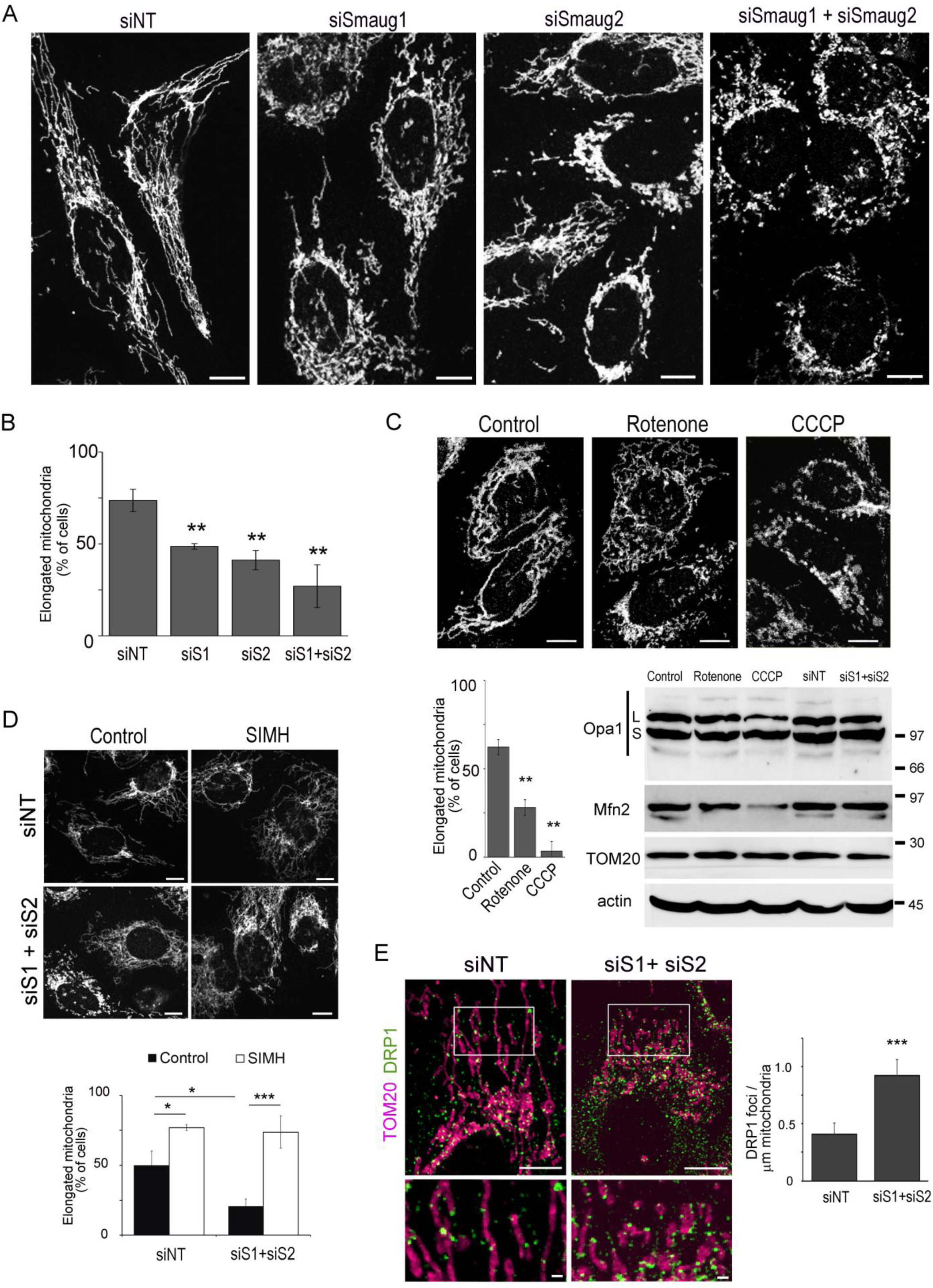
Mitochondrial network disruption upon Smaug1 and Smaug2 knockdown. **(A)** U2OS cells were treated with the indicated siRNAs and immunostained for TOM20. At least 200 cells per treatment were analyzed and the percentage of cells with elongated mitochondria is plotted **(B)**. A representative experiment out of three is shown. Scale bars, 10 μm. Error bars, standard deviation. **(C)** Cells were exposed to 2 µg/ml rotenone or 20µM CCCP for 1h or treated with the indicated siRNAs and whole protein extracts were analyzed by western blot for Opa1 and Mfn2. Opa1 L and S fragments are indicated. TOM20 and actin were analyzed as controls for mitochondrial content and protein loading respectively. In parallel, cells were immunostained for TOM20 and the fraction of cells with elongated mitochondria was counted in 200 cells from duplicate coverslips. **(D)** SIMH was induced as described in Materials and Methods and mitochondrial hyper-fusion was evaluated in at least 100 cells from duplicate coverslips for each experimental point. Means ± SD from two independent experiments are plotted (*, P < 0.05). Scale bars, 10 μm. **(E)** TOM20 (magenta) and DRP1 (green) were immunostained and the number of DRP1 puncta per 1 μm mitochondria were determined in 5-10 ROIs from 40 cells for each treatment. Means ± SD from two independent experiments are plotted (*, P < 0.05). Scale bars: whole cells, 10 μm, insets, 1 μm.

The mitochondrial network morphology depends on the balance between fusion and fission and we analyzed changes in molecular markers of both processes. Mitochondrial fusion requires Opa1 mitochondrial dynamin like GTPase (Opa1), mitofusin 1 (Mfn1) and mitofusin 2 (Mfn2). Reduced fusion is linked to changes in Opa1 protein cleavage and lower mitofusins levels. We found that both Opa1 and Mfn2 remained unaltered upon Smaug1 and Smaug2 double KD (Figure 3C). In contrast and as expected, CCCP elicited a strong response. Levels of Opa1 L fragment relative to S fragment levels were reduced, Mfn2 levels were downregulated, and all this correlated with decreased membrane potential (Figure 2D-E) and strong mitochondrial network fragmentation (Figure 3C) (Toyama et al., 2016). In addition, Opa1 and Mfn2 remained unaltered upon exposure to rotenone, which inhibits Complex I thus significantly triggering mitochondrial network fragmentation (Figure 3C, Toyama et al., 2016).

To further asses the integrity of the fusion machinery, we assessed a specific response termed stress-induced mitochondrial hyperfusion (SIMH) –which depends on Opa1 and Mfn1–, as previously described (Ehses et al., 2009; Tondera et al., 2009; see Methods). We found that SIMH was not altered upon Smaug1+2 KD. The fraction of siRNA NT-treated cells that showed elongated mitochondria increased from 50% to 77% upon SIMH induction. Smaug1+2 KD cells also responded significantly and the proportion of cells with elongated mitochondria increased from 21% to 74% after SIMH induction (Figure 3D). Collectively, these observations indicate that the mitochondria fusion machinery is fully functional in Smaug1+2 KD cells.

Mitochondrial fission is actively directed by recruitment of dynamin 1 like (DRP1) to the mitochondrial surface (Hoppins et al., 2007; Pitts et al., 1999) and we found that the number of DRP1 puncta recruited to mitochondria upon Smaug1+2 KD doubled the basal levels (4 vs 9 puncta per 10µm; Figure 3E). Altogether with the above results, these observations suggest that Smaug1+2 double KD activate mitochondrial fission likely as a consequence of defective mitochondrial function. Then, we used the mitochondrial network complexity as a read-out to further investigate the relevance of Smaug MLO formation.

First, we analyzed whether the mitochondrial phenotype provoked by Smaug1 and Smaug2 double KD is efficiently rescued by transfection of Smaug1 (Figure 4). We found that this is the case and in addition, the splicing variant ΔEIII, which has similar RNA binding and repression capacity (Figure 1 and Fernandez-Alvarez et al., 2016), also rescued the mitochondrial phenotype (Figure 4). Altogether, these observations suggest that the three mammalian Smaug isoforms reported to date, Smaug1; Smaug1 ΔEIII and Smaug2 play a rather redundant role.

**Figure 4.**
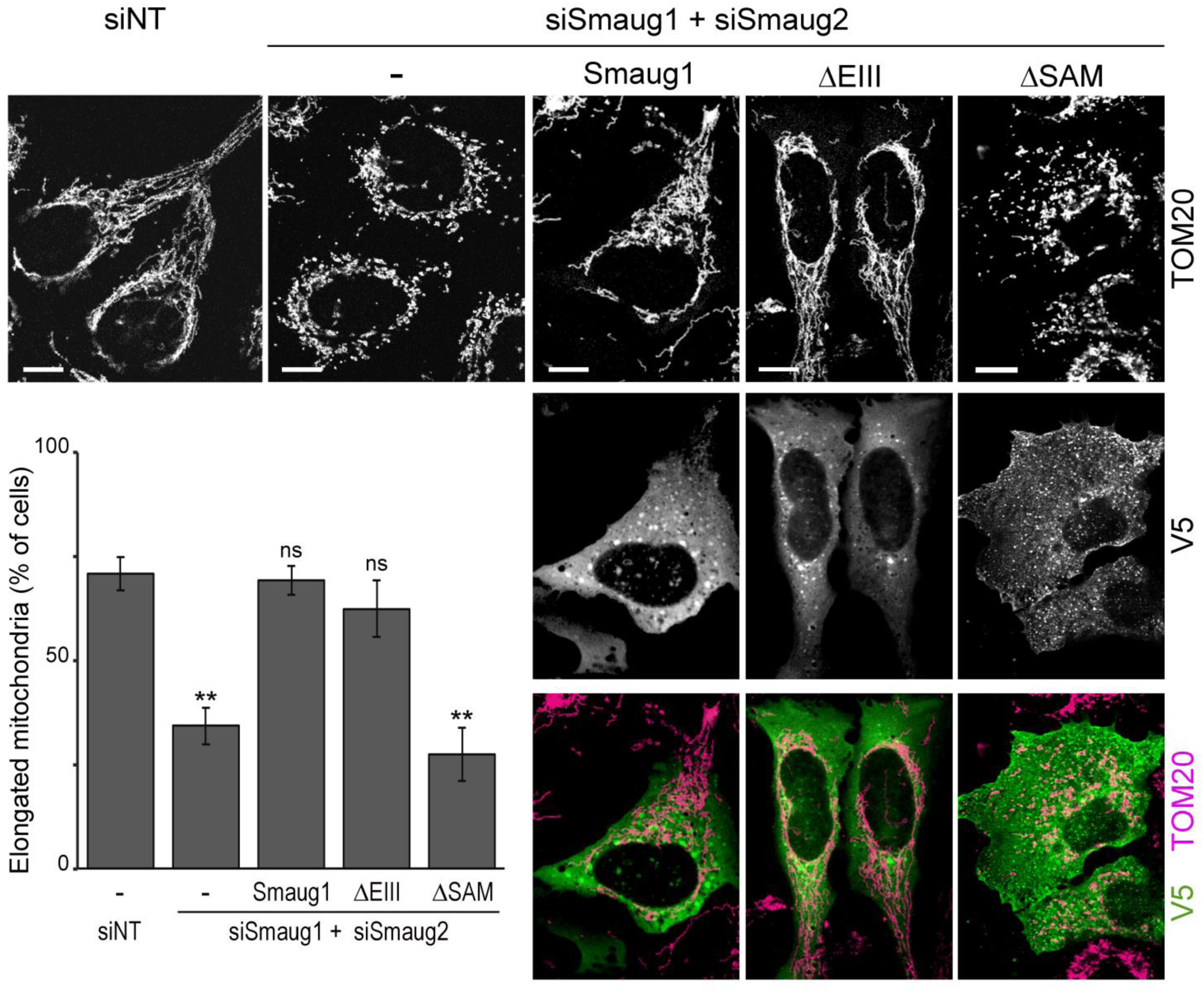
Smaug binding to RNA affects mitochondria. Smaug1, Smaug1 ΔEIII or Smaug1 ΔSAM tagged with V5-SBP were transfected simultaneously with siRNAs against Smaug1 and Smaug2. Cells were immunostained for TOM20 (magenta) and V5 (green). The fraction of cells with elongated mitochondria was determined in at least 200 cells from duplicate coverslips for each condition. A representative experiment out of three is shown. Scale bars, 10 μm. Error bars, standard deviation.

Smaug’s main function is to control mRNAs and we investigated the effect of a Smaug1 construct lacking the SAM domain and downstream region (termed ΔSAM), which are required for RNA binding (Aviv et al., 2003; Baez et al., 2011; Green et al., 2003; Johnson & Donaldson, 2006; Oberstrass et al., 2006). In three independent experiments, where more than 400 cells were analyzed for each construct, we found that deletion of the domain involved in RNA binding abrogates the rescue of the phenotype (Figure 4), strongly suggesting that mRNA regulation by Smaug1 is key to mitochondrial physiology.

### Condensation-defective Smaug1 mutants affect mitochondria

We aimed to investigate the relevance of Smaug1 MLOs condensation. First, we generated Smaug1 deletion mutants with defective body formation. Then, we used these tools as described in the next section to investigate the consequences of the lack of Smaug1-bodies on the mitochondrial network.

The domain organization of mammalian Smaug1 is depicted in figure 5A. Besides the SAM domain, mammalian Smaug orthologs include two different protein regions termed Smaug Similarity Region 1 (SSR1) and Smaug Similarity Region 2 (SSR2), which are conserved in flies. The yeast ortholog Vts1p includes only SSR1, which was shown to dimerize in vitro (Tang et al., 2007). We conducted transient-transfection experiments with the deletion constructs depicted in Figure 5A to analyze the formation of cytosolic condensates (Figure 5B-D). Similar expression levels were confirmed by immunoblot (Figure 5B). We found that deletion of the amino terminal region including SSR1 and SSR2 abrogates Smaug1-body formation. The ΔSSR1/2 fragment was always uniformly distributed in the cytosol in U2OS cells either fused to ECFP, V5 or V5-SBP (Figure 5C-D), as well as in HeLa and Cos-7 cells (data not shown). The SSR1-SSR2 fragment (SSR1/2) was required but was not sufficient to phase-separate as it always showed a uniform distribution (Figure 5C). Additionally, a construct that lacks the first 53 amino acids and termed ΔSSR1 was not able to form cytosolic bodies (Figure 5C-D). Finally, the splicing variant ΔEIII also formed cytosolic bodies (Figure 5C-D) as reported before (Fernandez-Alvarez et al., 2016).

**Figure 5.**
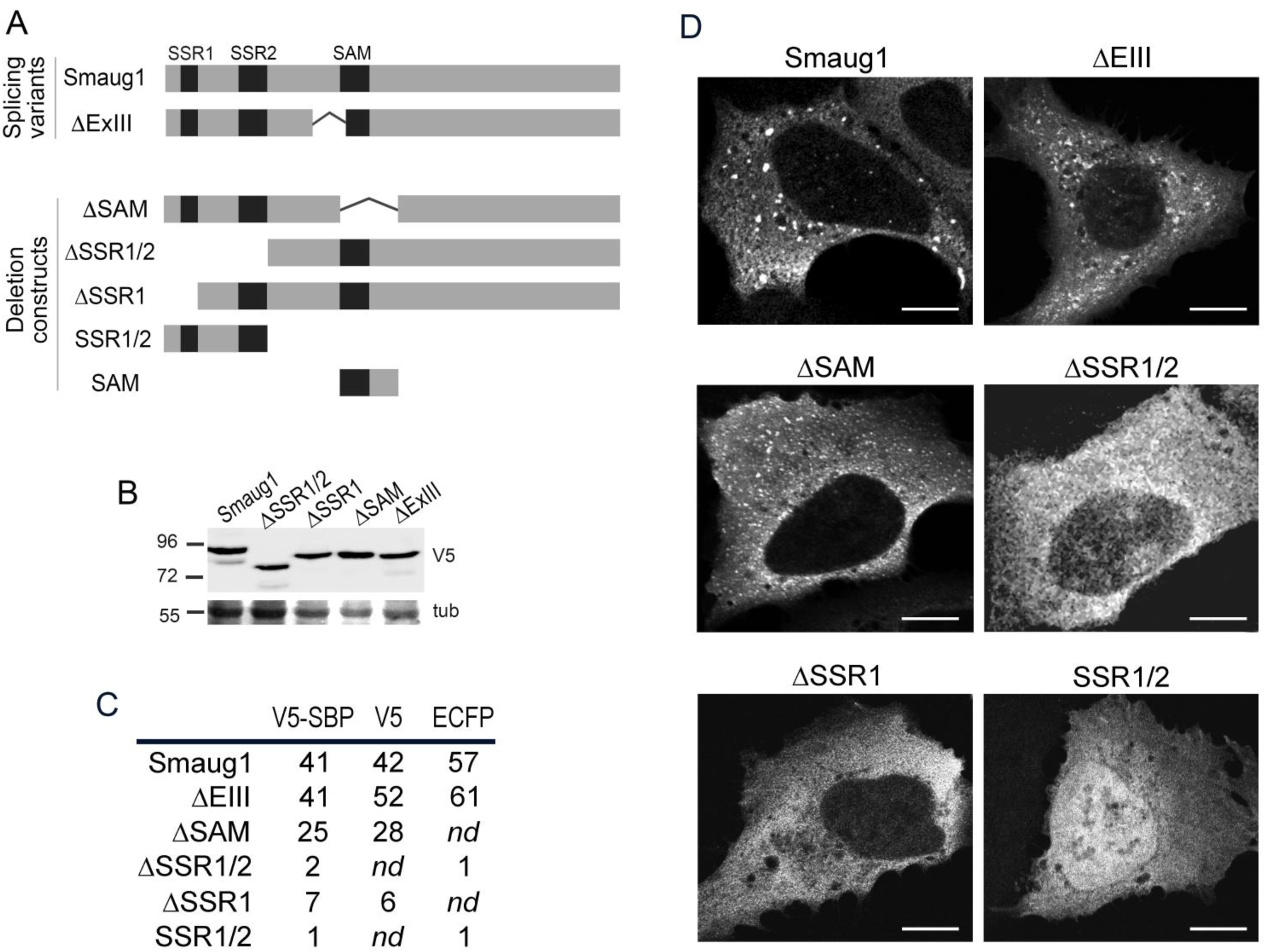
A conserved region required for Smaug1-body formation. **(A)** Human Smaug1 splicing variants and deletion constructs analyzed in this work. **(B-D)** The indicated constructs tagged with V5-SBP were transfected in U2OS cells and analyzed by western blot **(B)** and immunofluorescence **(C-D)**. Additional constructs tagged with V5 or ECFP were analyzed in independent experiments and the percentage of cells with cytosolic puncta was assessed by confocal microscopy. Duplicate coverslips and a minimum of 200 cells were analyzed in each case. Scale bars, 10 μm.

We next evaluated the recruitment of specific Smaug1 protein regions to the bodies formed by the full-length molecule. We co-transfected specific ECFP-tagged Smaug1 fragments with full length Smaug1 tagged with EYFP and measured the intensity of the ECFP tag inside and outside the Smaug1-EYFP bodies (Figure 6A). We found that SSR1/2-ECFP and the complementary construct ΔSSR1/2-ECFP, both of which remained uniformly distributed when expressed alone (Figure 5C), were efficiently recruited to the Smaug1-EYFP bodies. In contrast, the construct termed SAM-ECFP, which includes the SAM domain and 47 amino acids downstream, behaved similar to ECFP, both remained uniform and were not recruited to the Smaug1-EYFP bodies (Figure 6A).

**Figure 6.**
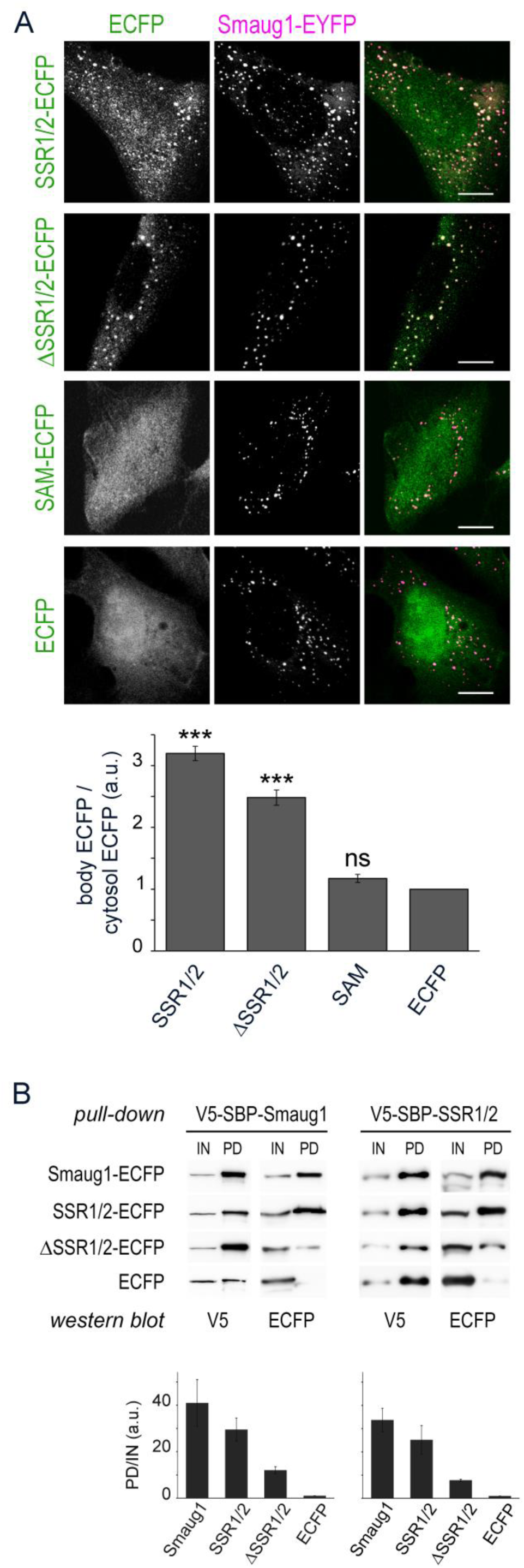
Protein-protein interactions mediate Smaug1-body formation. **(A)** Recruitment of the indicated Smaug1 fragments to Smaug1-EYFP bodies. U2OS cells were co-transfected with Smaug1-EYFP and the indicated ECFP-tagged constructs. Scale bars, 10 µm. ECFP signal intensity inside Smaug1-EYFP bodies relative to ECFP signal outside the bodies was measured in more than 150 Smaug1-bodies from randomly selected cells, and averaged values are plotted. Error bars, standard deviation. **(B)** V5-SBP-Smaug1 or V5-SBP-SSR1/2 was co-transfected with ECFP; Smaug1-ECFP; SSR1/2-ECFP; or ΔSSR1/2-ECFP. After pull-down with streptavidin-sepharose beads, the presence of V5- and ECFP-tagged constructs in cell lysates (IN) or pulled-down material (PD) was analyzed by western blot. A representative experiment out of three is depicted, and ECFP signal in the PD normalized to ECFP signal in the lysate is plotted for each construct. Error bars, standard deviation.

In parallel, we analysed physical interactions between these specific Smaug1 protein regions by pull-down assays. We co-expressed the Smaug1 fragments tagged with ECFP together with a full-length Smaug1 fused to a V5-SBP double tag. As expected, the pull-down of V5-SBP-Smaug1 efficiently recovered Smaug1-ECFP and did not significantly recover ECFP, which we chose as a negative control (Figure 6B, left). In addition, we found that the SSR1/2-ECFP fragment co-purified with V5-SBP-Smaug1, whereas the ΔSSR1/2-ECFP construct was recovered less efficiently (Figure 6B, left). Conversely, V5-SBP-SSR1/2 significantly co-pulled down Smaug1-ECFP and SSR1/2-ECFP (Figure 6B, right). In contrast, the ΔSSR1/2-ECFP construct was recovered less significantly and ECFP was almost absent from the material co-pulled down with V5-SBP-SSR1/2. In all cases, a contribution of endogenous Smaug1 and Smaug2 that may bridge the transfected Smaug1 fragments together cannot be ruled out.

Despite minor quantitative differences with the outcome of the imaging analysis described above –likely due to the harsher conditions of pull-down assays–, altogether these observations suggest that Smaug1 phase separation is driven by multiple protein regions. It likely involves SSR1 homodimerization and additional interactions between unknown protein motifs distributed along the molecule.

We found that RNA regulation by Smaug1 affects mitochondrial function (Figure 4), and we wondered whether condensation of Smaug1-bodies is relevant as well. We found that truncated Smaug1 constructs that show defective body formation failed to rescue the mitochondrial phenotype (Figure 7). In three independent experiments, cells treated with siRNAs against Smaug1 and Smaug2 and transfected with V5-SBP-ΔSSR1/2 or V5-SBP-ΔSSR1 showed mostly short mitochondria, with values similar to those of cells transfected with V5-SBP-ΔSAM which, as seen in figure 4, did not rescue the phenotype. In contrast, transfection of full-length Smaug1 rescued the mitochondrial network complexity in most cells, as previously seen. We conclude that mitochondria depend on both RNA binding by Smaug and Smaug-body formation.

**Figure 7.**
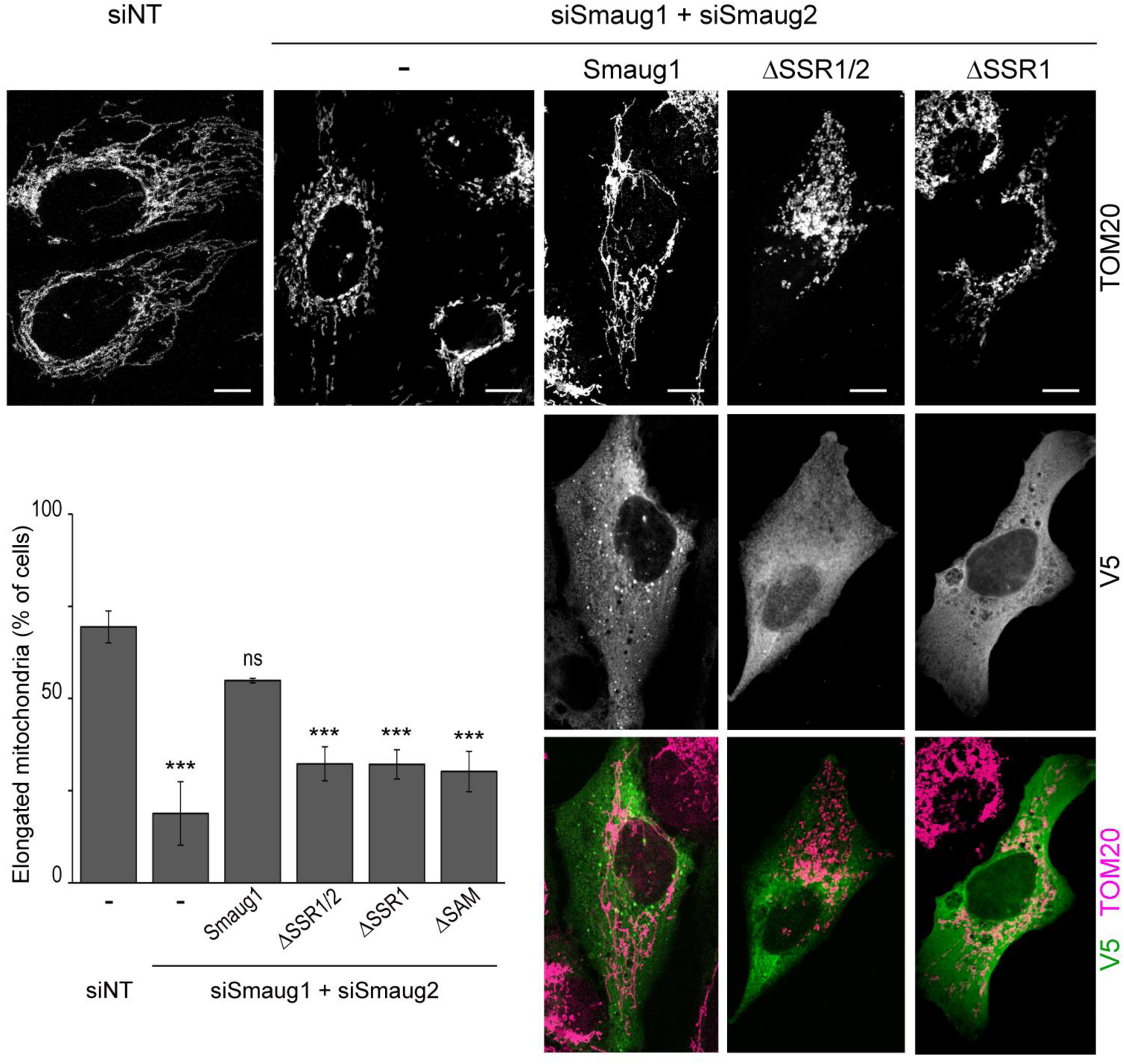
Defective Smaug1-body formation correlates with mitochondrial defects. U2OS cells were treated with non-targeting siRNA (siNT) or with siRNAs against Smaug1 and Smaug2 and transfected with the indicated constructs tagged with V5-SBP (green). The mitochondrial network was analyzed by TOM20 immunostaining (magenta) in at least 200 cells from duplicate coverslips. A representative experiment out of three is shown. Scale bars, 10 µm. Error bars, standard deviation.

### Smaug1 MLO formation affects RNA binding

We next investigated the consequences of defective Smaug1-body formation on RNA regulation. In order to uncouple RNA binding from translational repression, we used a MS2-tethering strategy, as previously described (Bhandari et al., 2014; Deng et al., 2015). Briefly, a firefly luciferase reporter carrying 6 MS2 binding sites (6xMS2bs) or a control reporter with no MS2 binding sites were co-transfected with the above described Smaug1 fragments fused to a double tag MS2-HA, to direct its tethering to the firefly luciferase reporter and to allow its detection by western blots, respectively. In six independent experiments, we found that Smaug1 reduced firefly luciferase 6xMS2bs expression to about 50% of the control levels (Figure 8A). Deletion of SSR1 and SSR2 repressed the reporter to the same extent (ΔSSR1/2, Figure 8A). As expected, tethering of the ΔSAM construct repressed the reporter similarly to the full length Smaug1, while tethering the SAM domain had no effect (Figure 8A). For comparison, the effect of SMG7 –a key factor of the non-sense mediated decay pathway– was analyzed and found to strongly repress the luciferase reporter (Figure 8A). RNA levels were not affected by tethering of either full length or truncated Smaug1 constructs, whereas SMG7 induced a significant decay (51±4% relative to the MS2 control). Similar results were observed in HEK293T (Figure 8A) and U2OS cells, where Smaug1-MS2-HA repressed the reporter to 55±5%; ΔSSR1-MS2-HA, to 57±10% and ΔSSR1/2-MS2-HA, to 48±6%, whereas tethering of the SAM domain elicited no effect.

**Figure 8.**
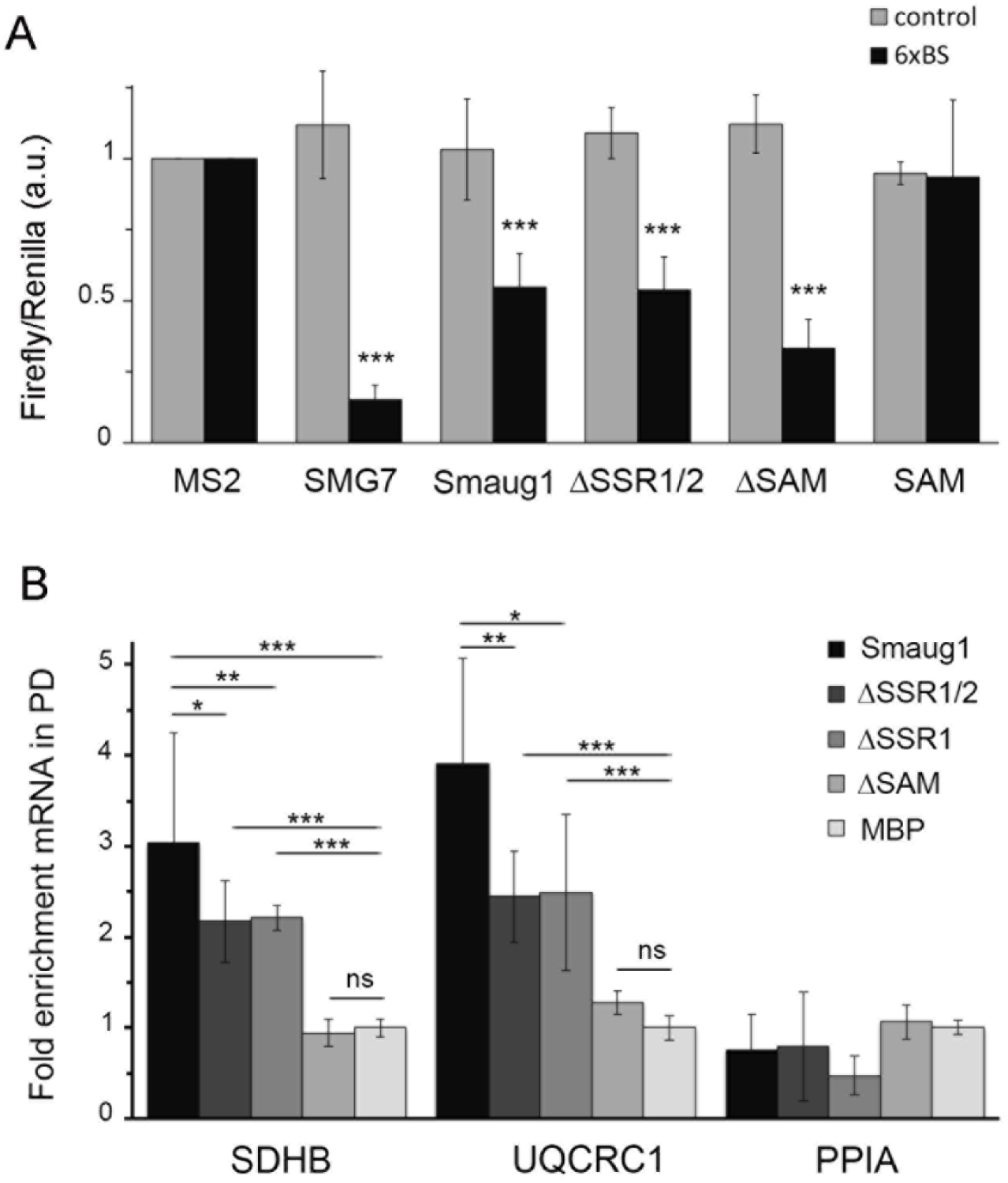
Smaug1 condensation affects mRNA binding. **(A)** Tethering assay of the indicated Smaug1 constructs tagged with MS2-HA. Two firefly reporters –with or without a tandem array of 6 MS2 binding sites (6xBS)– were analyzed in parallel and normalized to a co-transfected renilla reporter. The firefly/renilla ratio averaged from six independent experiments is plotted. Error bars, standard deviation. **(B)** RNA pull-down. The indicated constructs tagged with V5-SBP were pulled-down and the recovery of the indicated mRNAs was determined by qRT-PCR as in Figure 1. Error bars, standard deviation.

Theoretically, the 6 MS2 binding sites present in the firefly luciferase reporter may tether up to 6 protein molecules, and we considered whether this may drive the phase separation of the otherwise “soluble” Smaug1 deletion constructs. Confocal imaging analysis indicated that less than 10% of the cells co-transfected with ΔSSR1/2-MS2-HA and the 6xMS2bs reporter showed microscopically visible puncta after staining of HA, whereas, as expected, more than 50% of the cells co-transfected with Smaug1-MS2-HA and the 6xMS2bs reporter showed Smaug1-MS2-HA bodies. The presence of nanoscale condensates cannot be ruled. Collectively, these observations indicate that the formation of Smaug1-bodies does not significantly affect mRNA repression when Smaug1 is strongly tethered to multiple sites in the target transcript.

Next, we investigated whether Smaug1 phase separation affects the interaction with endogenous mRNAs. We focused on SDHB and UQCRC1 mRNAs, which associate to Smaug1 MLOs and co-purified with Smaug1 (Figure 1). As expected, whereas SDHB and UQCRC1 mRNAs showed a three-fold enrichment in the Smaug1 pull-down, the construct lacking the SAM domain failed to bind to these transcripts (Figure 8B). Relevantly, we found that deletion of either the SSR1 or the N-terminal region including SSR1 and SSR2 moderately impaired the recovery of SDHB and UQCRC1 mRNAs. In several independent experiments, the pull-down of condensation-defective V5-SBP-ΔSSR1 or V5-SBP-ΔSSR1/2 (Figure 5C) recovered 60-75% of the amount of mRNA that co-purified with the full-length construct (Figure 8B). In all cases, we found that the amount of SDHB and UQCRC1 mRNAs in the whole extract remained unaltered. Expression levels and pull-down recovery of the V5-SBP-tagged constructs were confirmed to be quantitatively similar by western blot (Supplementary file 4). These results indicate that the Smaug1-body formation helps –but it is not strictly required– for the interaction of Smaug1 with its target mRNAs.

### Mitochondrial activity affects Smaug1-body condensation

The above observations suggest that the condensation of Smaug1-bodies contributes to the regulation of mitochondrial relevant mRNAs. We further speculated that Smaug1-bodies may respond to changes in mitochondrial activity, which we manipulated by pharmacological approaches. We assessed the effect of complex I inhibition by rotenone on Smaug1-bodies detected by staining of the endogenous molecule. We found that exposure to rotenone for one hour induced the dissolution of the Smaug1-bodies in a significant proportion of cells. On average, the number of cells with Smaug1-bodies dropped to 66% the basal values. Smaug1-body disappearance did not correlate with mitochondrial fragmentation (Figure 9A). Smaug1-EYFP bodies were similarly affected and we found that the proportion of cells with Smaug1-EYFP bodies was reduced from 66 % to 42 % after 1h exposure to rotenone. In contrast, strong depolarization by CCCP elicited no effect on Smaug1-EYFP bodies (Figure 9B), while seriously affecting the mitochondrial network (Figure 2C-D and Figure 3B). These observations indicate that Smaug1-body disappearance does not correlate with mitochondrial damage or fragmentation but rather with complex I inhibition.

**Figure 9.**
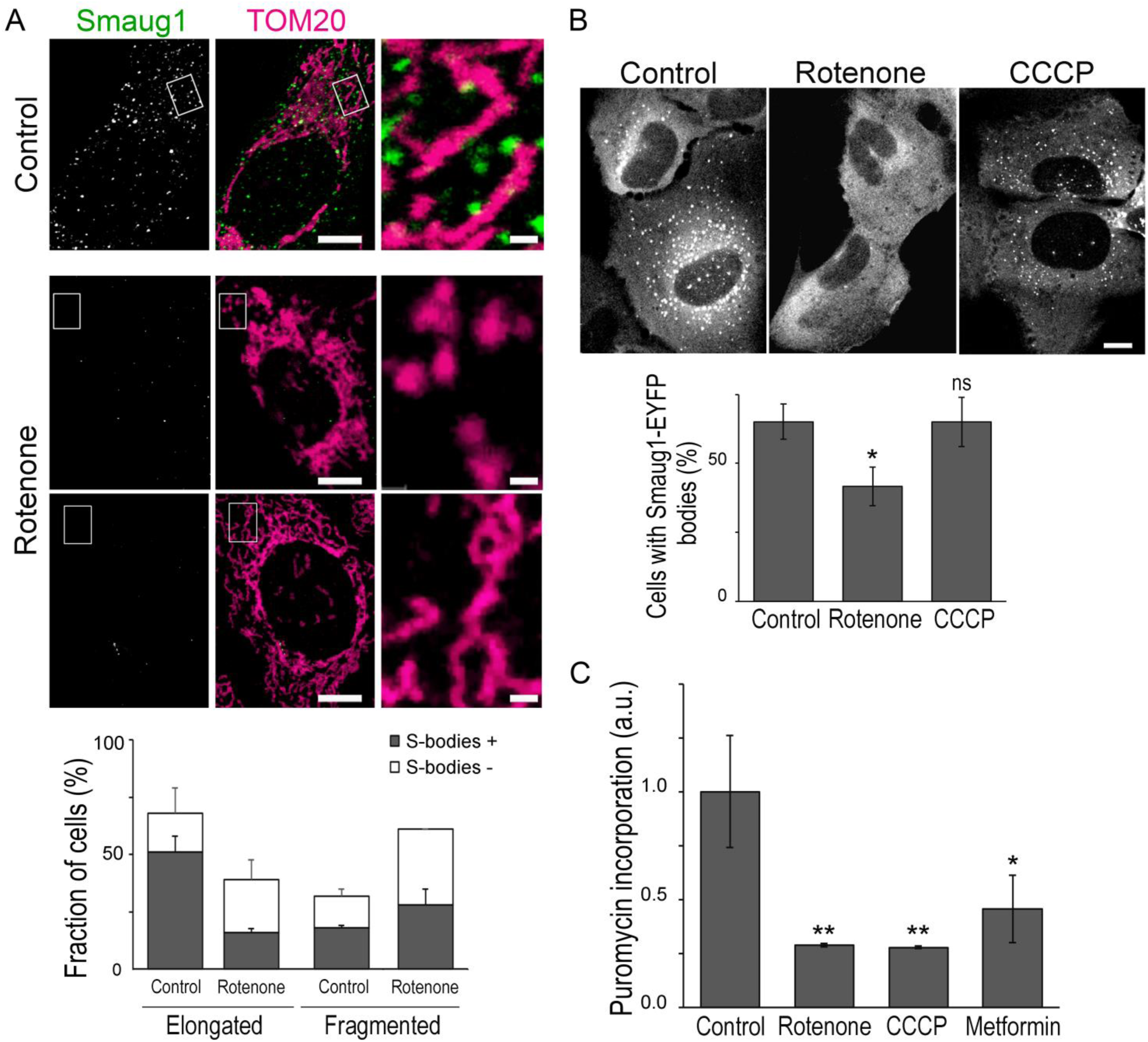
Rotenone induces Smaug1-body dissolution. **(A)** U2OS cells were exposed to 2µg/ml rotenone for 1 hour and stained for Smaug1 (green) and TOM20 (magenta). Two representative cells showing Smaug1-body disappearance upon rotenone with either elongated (upper panel) or fragmented mitochondria (lower panel) are depicted. The presence of Smaug1-bodies was evaluated in at least 200 cells from duplicate coverslips for each condition as described in Material and Methods. Scale bars: whole cells, 10 μm, zooms, 1 μm. Error bars, standard deviation. **(B)** Cells transfected with Smaug1-EYFP were treated during 1 h with 2 µg/ml rotenone or 20 µM CCCP and analyzed by confocal microscopy. More than 200 cells in duplicate coverslips were analyzed for each experimental point. Scale bars, 10 µm. Error bars, standard deviation. **(C)** Cells were exposed to 2 µg/ml rotenone, 20 µM CCCP or 100 µM metformin for one hour and immediately incubated with puromycin during 5 min. Puromycin incorporation was analyzed with specific antibodies as described in Materials and Methods in at least 80 cells for each experimental point.

Dissolution of PBs, SGs, Smaug1-bodies and related MLOs can be the consequence of translation activation. We analyzed the effect of rotenone and CCCP in the global translation rate, which we measured by puromycin incorporation. Briefly, cells were exposed to a short pulse (5 min) of puromycin, which was then detected by immunofluorescence. As expected, rotenone and CCCP seriously reduced translation (Fessler et al., 2020) (Figure 9C). These observations indicated that Smaug1-body dissolution does not correlate with changes in global translation.

Next, we investigated the effect of metformin, a drug highly relevant in the clinics that regulates the energetic metabolism through multiple pathways including inhibition of mitochondrial complex I (Hawley et al., 2002; Wheaton et al., 2014). We found that the number of cells with Smaug1-bodies detected by staining of endogenous Smaug1 decreased to about 2/3 of the basal values after 1-hour treatment with 100 µM metformin (Figure 10 A). In accordance with previous observations (Izzo et al., 2017; Wang et al., 2017) metformin elicited no effect on the mitochondrial network. In addition, we found that metformin moderately reduced puromycin incorporation (Figure 9C), suggesting that translation rate is slightly downregulated, as reported previously (Larsson et al., 2012).

**Figure 10.**
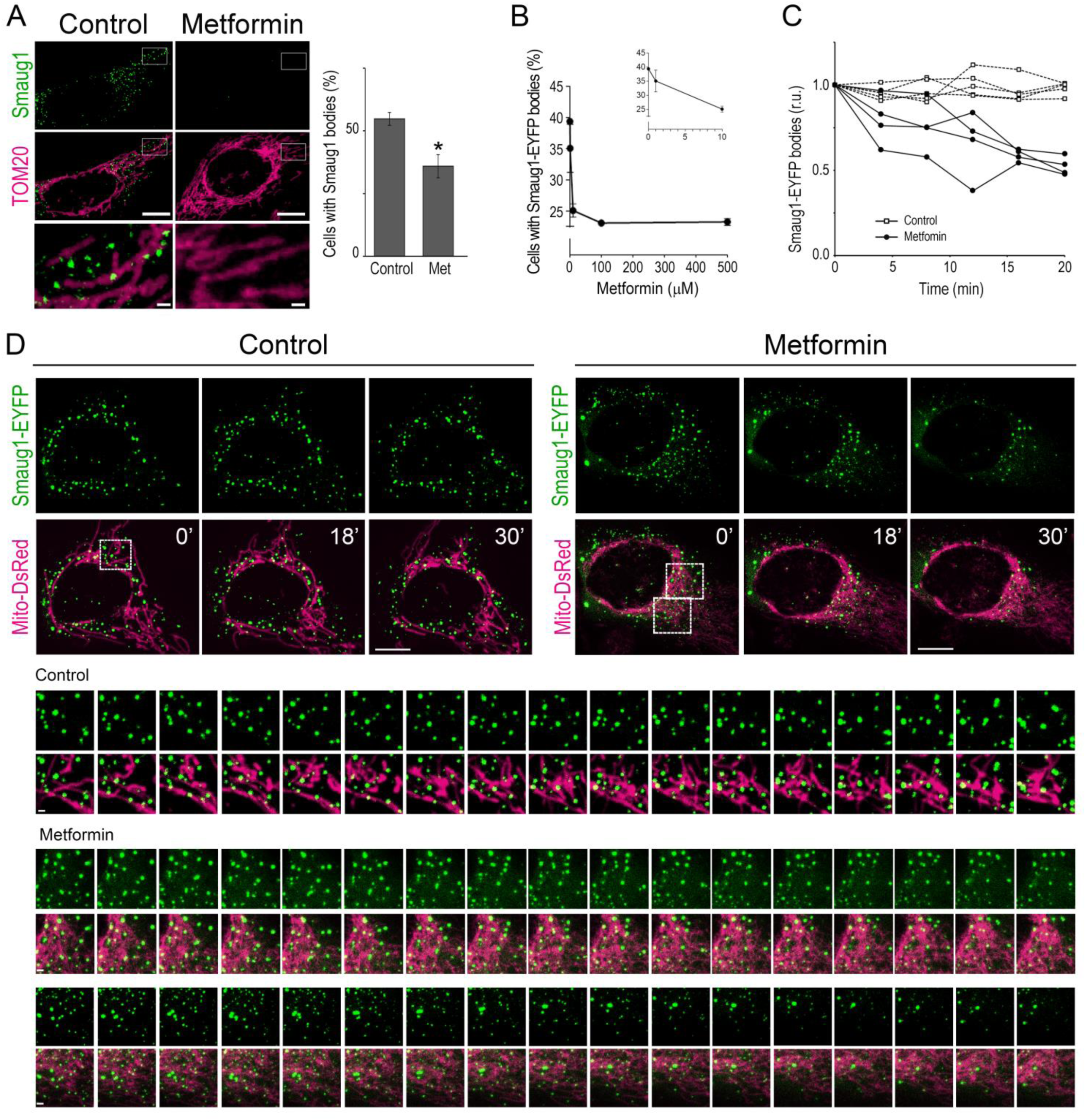
Metformin induces Smaug1-body dissolution. **(A)** U2OS cells were exposed to 100 µM metformin (Met) and stained for Smaug1 (green) and TOM20 (magenta). The presence of Smaug1-bodies was evaluated as in Figure 9. **(B)** Cells transfected with Smaug1-EYFP were exposed during 1 h at the indicated metformin concentrations. More than 100 cells from duplicate coverslips were analyzed for each experimental point. Error bars, standard deviation. **(C-D)** Live cell imaging of U2OS cells transfected with Smaug1-EYFP and Mito-DsRed and exposed to 100 µM metformin. **(C)** Smaug1-EYFP body number from four representative control and four metformin-treated cells normalized to pre-treatment values as described in Material and Methods are plotted. **(D)** A representative control cell and a metformin-treated cell are shown at 0, 18, and 32 min after treatment initiation. Magnifications of the indicated areas with 2-min intervals from 0 to 32 min are shown. Scale bars, whole cells, 10 μm, zooms, 1 μm.

Smaug1-EYFP bodies responded similarly to metformin in a dose-dependent manner (Figure 10B). The number of cells with Smaug1-EYFP bodies was reduced to half of the control values after 1-hour exposure to 100 µM metformin (Figure 10B). Time-lapse confocal microscopy of Smaug1-EYFP transfected cells showed that the response is fast. As in fixed cells, we found that about half of the cells responded to metformin exposure and in responsive cells Smaug1-EYFP bodies started to dissolve immediately after treatment. Smaug1-EYFP bodies showed a gradual reduction in size and /or fluorescence intensity and a significant proportion of bodies completely vanished (Figure 10C-D, Video1-2).

We have previously shown that synapse activation can induce Smaug1-body dissolution and the release of specific mRNAs together with a global translational shutdown (Baez et al., 2011; Luchelli et al., 2015). We speculated that similarly, Smaug1-body dissolution upon exposure to metformin is linked to the release of mRNAs bound to Smaug1. We performed co-pulldown assays as before (Figure 1) and found that the exposure to metformin reduced the amount of SDHB mRNA that co-pulled down with V5-SBP-Smaug1, whereas SDHB mRNA total levels remained unchanged (Figure 11A-B). Binding of UQCRC1 mRNA was affected to a lesser extent. Collectively, these observations indicate that metformin triggers Smaug1-body dissolution and the release of associated mRNAs. Whether released transcripts are then engaged in translation remains to be investigated.

**Figure 11.**
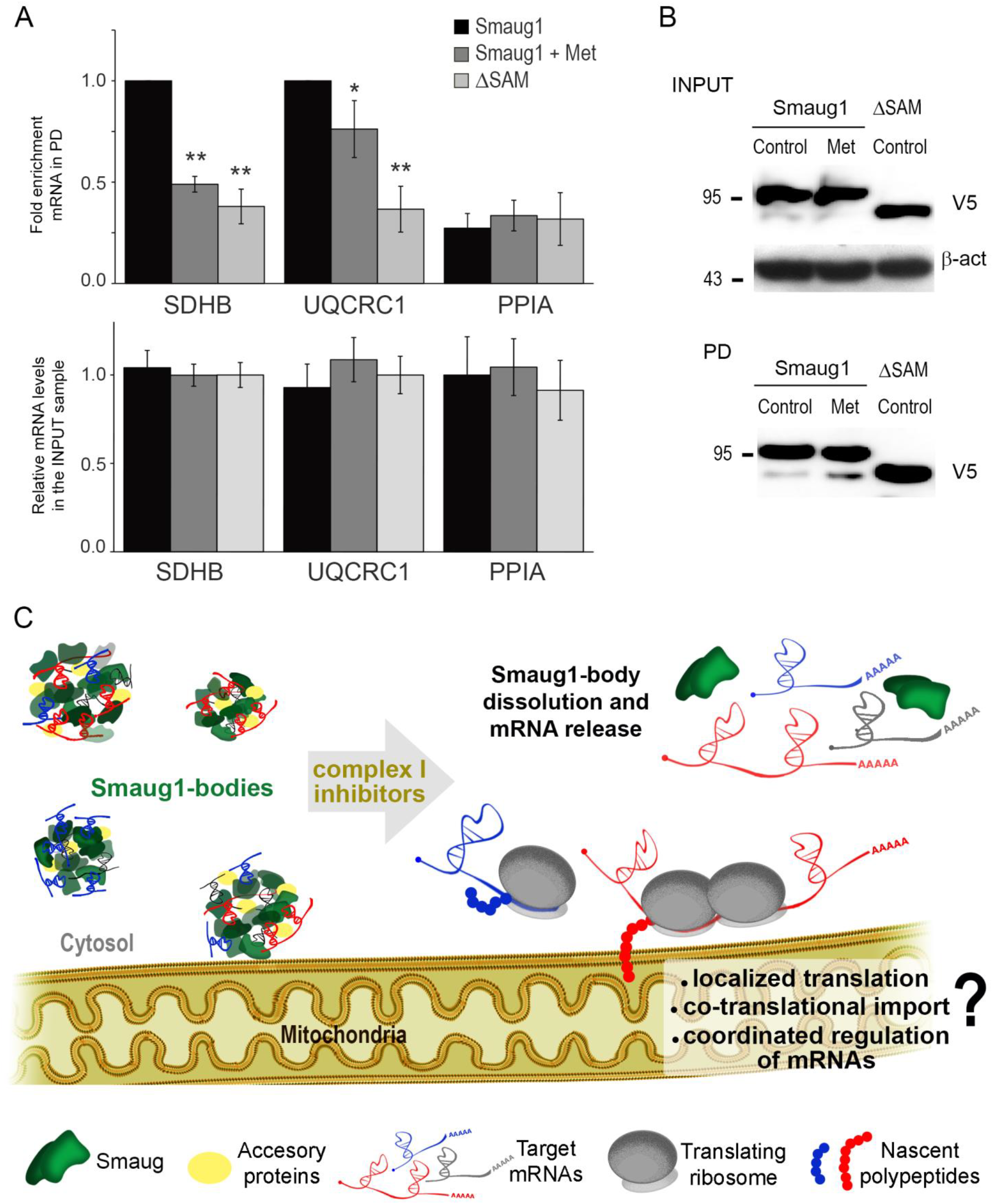
Metformin reduces the interaction with target mRNAs. **(A-B)** U2OS cells transfected with V5-SBP-Smaug1 were exposed to 100 µM metformin for one hour and the co-pulldown of SDHB, UQCRC1 and PPIA mRNAs was determined by qRT-PCR as in Figure 1. Cells transfected with V5-SBP-ΔSAM were analyzed in parallel. **(A)** The pull-down/input ratio (upper panel) and total levels in the input samples (bottom panel), both normalized to RpLp0 mRNA values, are plotted for each transcript. Error bars, standard deviation. **(B)** Western blot of the input and pull-down (PD) samples revealed with anti-V5 antibody. **(C)** Proposed model. Protein-protein interactions including SSR1 homodimerization drive Smaug1-body formation (green). Additional proteins (yellow) are expected to be present in Smaug1-bodies and whether Smaug2-bodies are separate MLOs is currently unclear. Smaug1-bodies contain mRNAs that encode mitochondrial proteins, including SDHB and UQCRC1 and likely many others. Changes in mitochondrial activity triggers Smaug1-body dissolution and mRNA release, likely enabling their coordinated translation, which may occur at the mitochondrial periphery.

## Discussion

Here we report that defective Smaug1 MLO condensation correlates with altered mitochondrial function. In addition, Smaug1 MLOs contain SDHB and UQCRC1 mRNAs and exposure to mitochondrial inhibitors rapidly triggers Smaug1-body dissolution and mRNA release. We hypothesize that dynamic Smaug1-body formation links mitochondrial function with the regulation of mRNAs that encode mitochondrial enzymes (Figure 11C).

As reported before in neurons, we found that Smaug1-bodies in U2OS cells exclude the translational apparatus and dissolve when mRNAs are trapped into polysomes –as exemplified by cycloheximide exposure (Figure 1)–, supporting the notion that Smaug1-bodies contain repressed mRNAs that can be released to enter translation. Smaug1-bodies are different from PBs, which coordinate the expression of sets of mRNAs functionally related and termed “RNA regulons” (Hubstenberger et al., 2017). In addition to UQCRC1 and SDHB mRNAs a number of transcripts that code for mitochondrial enzymes were reported to bind Smaug orthologs, thus defining a mitochondrial regulon linked to Smaug-bodies (Chartier et al., 2015; Chen, Dumelie, et al., 2014; Tadros et al., 2007). The collective dysregulation of SDHB, UQCRC1 and additional mRNAs is likely the direct cause of the serious mitochondrial phenotype provoked by Smaug1 loss of function, characterized by altered OXPHOS activity and disrupted mitochondrial network.

Current models for MLO formation propose that multiple protein-protein interactions together with protein-RNA and RNA-RNA contacts direct a LLPS process which eventually drive the condensation of ribonucleoparticles in specific bodies (Courchaine et al., 2016; Guo & Shorter, 2015; Perez-Pepe et al., 2018; Sachdev et al., 2019; Van Treeck et al., 2018). MLOs involved in mRNA regulation can be complex in composition and several factors that interact with Smaug orthologs, including Argonaute, 4E-binding proteins and deadenylase complexes might additionally help Smaug1-body formation (Amadei et al., 2015; Chartier et al., 2015; Nelson et al., 2004; Pinder & Smibert, 2013). We were able to identify conserved Smaug1 domains involved in Smaug1-body formation. We propose that SSR1 dimerization and additional interactions involving further protein regions act together with RNA binding to direct Smaug1 phase separation. We found that Smaug1 condensation does not directly influence translation repression but affects the interaction with target mRNAs (Figure 8). Among several mechanisms, the presence of multiple RNA-binding domains in the Smaug1-body will increase the avidity for mRNA molecules. A similar model was proposed for the yeast ortholog Vts1 (Bieman, 2014).

We found that defective Smaug1-body condensation elicits a moderate effect in RNA binding while mitochondria are severely affected (Figure 7). We propose that the mitochondrial defects are the consequence of concurrent mechanisms that remain to be further investigated. First, a single Smaug1-body may contain several mRNA species, as exemplified here by UQCRC1 and SDHB mRNAs, whose co-occurrence in Smaug1-bodies is higher than random (Figure 1F). This may enable the coordinated regulation of several target mRNAs. Second, Smaug1-bodies contact mitochondria and we speculate that this may help local regulation. Several nuclear messengers that encode mitochondrial proteins including complex I and II subunits are translated at the mitochondria periphery in yeast, plant and animal cells. Localized translation can enable co-translational import and assembly, thus helping the biogenesis of multi-subunit complexes (Gehrke et al., 2015; Schatton & Rugarli, 2018; St-Pierre & Topisirovic, 2016; Vincent et al., 2017; Wilk et al., 2016; Williams et al., 2014; Wu et al., 2018). Relevantly, biogenesis of succinate dehydrogenase complex involves a strict co-translational control with the participation of multiple co-factors, and mutations in assembly factors lead to serious mitochondrial defects (reviewed in (Moosavi et al., 2019)). This highlights the significance of the co-translational regulation of SDH and additional ETC complexes, many of them under Smaug control (Chartier et al., 2015; Chen, Dumelie, et al., 2014; Tadros et al., 2007). Smaug bodies may parallel the function of MLOs formed by the RNA binder Tis11, which associates to the ER thus influencing ER-associated translation (Bethune et al., 2019; Ma & Mayr, 2018). Our observations suggest that Smaug1+2 bodies may affect mitochondria-associated translation, by helping mRNA transport and/or allowing coordinated mRNA de-repression upon dissolution. These important hypotheses are worthy of further studies.

Translation of mitochondrial enzymes responds to mitochondrial physiology and energetic balance (Bethune et al., 2019; Gao et al., 2014; Schatton et al., 2017; Wakim et al., 2017; Wu et al., 2018). Relevantly, along with the significance of Smaug1-body formation to mitochondrial function, we found that acute complex I inhibition triggers Smaug1-body dissolution (Figure 9 and 10, and videos 2 and 3). We have recently shown that Drosophila Smaug MLOs are affected by Smoothened (Smo), a key molecule in the Hedgehog (HH) pathway (Bruzzone et al., 2020). Whether Smaug regulation by Smo is conserved in mammals and whether this may occur in connection with mitochondrial activity is currently unknown. Messenger RNA release from dissolving MLOs is linked to translation (Khong & Parker, 2018; Moon et al., 2019) and thus, we speculate that Smaug1-body dissolution allows the translation of SDHB, UQCRC1 and other enzymes to recover mitochondrial function. Remarkably, rotenone and metformin trigger Smaug1-body dissolution along with widespread translational repression, as similarly occurs in neurons exposed to specific neurotransmitters (Baez et al., 2011; Luchelli et al., 2015). Collectively, these observations suggest that Smaug1-bodies respond to specific cellular cues to release selected mRNAs, yet at the same time most messengers are massively repressed, thus providing a mechanism for selective translation (Thomas et al., 2014). To our knowledge, Smaug1 bodies are the first mRNA-containing MLO that can respond to mitochondrial activity.

The role of Smaug orthologues as post-transcriptional regulators of energetic metabolism appears evolutionary conserved as yeast Vts1p –which forms MLOs and prion-like condensates– controls mRNAs linked to nutrient sensing (Chakravarty et al., 2020; She et al., 2017). Translation of nuclear-encoded transcripts at the mitochondrial periphery is known to be regulated by a reduced number of RNA-binding proteins, including the highly conserved factors clustered mitochondria homolog (CLUH) and Pumilio orthologs (reviewed in(D’Amico et al., 2019; Gao et al., 2014; Gehrke et al., 2015; Kopp et al., 2019; Lee & Tu, 2015; Olivas & Parker, 2000; Pla-Martin et al., 2020; Schatton et al., 2017; Schatton & Rugarli, 2018; Vardi-Oknin & Arava, 2019; Wakim et al., 2017) The putative interplay between these pathways and Smaug remains to be investigated.

Besides its relevance to cancer and diabetes, mitochondrial activity is particularly important in normal tissues with high energy demands, including neurons and muscle cells. Mitochondrial defects are causative of several neurological conditions and muscular dystrophies (Friedman & Nunnari, 2014; Gan et al., 2018; Kriaucionis et al., 2006; Rangaraju et al., 2019; Rugarli & Langer, 2012; Shan et al., 2019). The strong effect of Smaug1 and 2 in neuronal development and maturation, and in muscular cells in connection with myopathies is likely related to the relevance of Smaug to mitochondrial function (Amadei et al., 2015; Baez et al., 2011; Chartier et al., 2015; Chen, Holland, et al., 2014; de Haro et al., 2013; Luchelli et al., 2015). The present work opens new questions on the motility and dynamics of Smaug-bodies in a wide diversity of cellular contexts. A major issue to be addressed is how these MLOs respond to metabolic cues and what are the consequences in the transport and translation of mRNAs that encode mitochondrial enzymes.

## Materials and Methods

### Cell culture and treatments

U2OS and HEK293T cells were obtained from the American Tissue Culture Collection (ATCC) and grown and maintained as indicated. Cell lines were transfected with Jet Prime (Polyplus Transfection) according to manufacturer’s instructions.

pCDNA6-V5-hSmaug1 (Smaug1-V5) and pECFP-N1-hSmaug1 (Smaug1-ECFP) constructs were previously described (Baez y Boccaccio 2005). Smaug1-EYFP was prepared by direct sub-cloning of pECFP-N1-hSmaug1 into pEYFP-N1. ΔSAM-hSmaug1-ECFP and ΔSAM-hSmaug1-EYFP with a deletion of amino acids 318 to 424 were constructed by SOE-PCR using ECFP-ΔC-hSmaug1 and hSmaug1-ECFP as templates and the following primers (SOE-1: 5’-GCGAGCCTCGGGTGTGTTACGAGC-3, SOE-2: 5’-GCTCGTAACACACCCGAGGCTCGC-3’). ΔSAM -hSmaug1-V5 was obtained by insertion into pcDNA6.0-B-His-V5 using HindIII/XhoI restriction digestion. ECFP-ΔC-hSmaug1 which comprises amino acids 2 to 418 was constructed by PCR using T7 forward and 5’AAGGCGAGCCTCGAGGGTG-3’ reverse primers and pCDNA6-V5-hSmaug1 as template with XhoI/HindIII deletion fragment inserted in a SalI/HindIII pECFP-C3 digested vector. ΔSSR1-hSmaug-V5 spans amino acids 54 to 718 and was constructed by PCR using forward primer 5’GCTCCCAAGCTTACCATGGAGCTGCACGTCCTCGAACG 3 ’ and reverse primer 5’ GCTCCGCTCGAGAAGATGGTGGAGGTCCGGTCAAC 3’ and hSmaug1-V5 as template and pcDNA6.0-B-His-V5 as vector. SSR1/2-hSmaug1-ECFP was constructed by insertion of a HindIII/SalI fragment into pECFP-N1 and spans amino acids 2 to 154. All MS2-HA constructs were obtained by PCR using primers carrying BamHI and XhoI restriction sites into the pcDNA3.1-MS2-HA (Lykke-Andersen et al., 2000). V5-SBP-hSmaug1 was cloned by PCR using primers carrying XhoI and BamHI sites into pT7-V5-SBP (Sgromo et al., 2017). V5-SBP-ΔSAM-hSmaug1 was sub-cloned by KpnI/BamHI restriction of ΔSAM-hSmaug1-ECFP. V5-SBP-ΔN-hSmaug1 was cloned by BamHI/SacII digestion into a BglII/SacII pT7-V5-SBP-C1 digested plasmid. V5-SBP-ΔN-hSmaug1 was cloned by BamHI/EcoRI digestion into a BglII/EcoRI pT7-V5-SBP-C1 digested plasmid.

For knock-down experiments, U2OS cells were treated with 50 nM siRNAs using Jet Prime transfection reagent for 48 hs following the manufacturer’s instructions. siRNAs were obtained from Dharmacon or Eurofins and were: NT, UAGCGACUAAACACAUCAA; Samd4A mix: 1-GACCAGAGGGUUUGCGAA, 2-CUACAGGUAUAUAGCUCAA, 3-CUUAAUGAAAUCCGAACAA, and 4-GAUGGAAAUGACAGCGCUA; Samd4B mix, or sequences 1 or 4 alone: 1-ACACAGAGGCCAAGUCGGA, 2-CAUGAGGCUUUCACGGAGA, 3-AUCCAGAAGCUGCGUGAGA, 4-GCUGAAGCUCCUCCGGACA.

Drugs treatments were as follows: metformin hydrochloride (Sigma-Aldrich) was dissolved in H_2_O and used during 1h at 500 µM, unless otherwise indicated. Rotenone was used at 2µg/ml (from a stock solution 50mg/ml in DMSO) during 1h unless otherwise indicated. Carbonyl cyanide m-chlorophenyl hydrazine (CCCP) was used 20 µM (from a stock solution 100 mM in DMSO) during 1 h unless otherwise indicated. To induce SIMH, cells were exposed to 1μM cycloheximide for 3 hours (Ehses et al., 2009; Tondera et al., 2009).

### Immunofluorescence, fluorescent in situ hybridization and puromycilation

Immunofluorescence of cultured cells was performed after fixation, permeabilization and blocking as usual (Baez et al., 2011; Fernandez-Alvarez et al., 2016; Luchelli et al., 2015). Primary antibodies were diluted as follows: V5 1:500, anti-SAMD4A HPA043061 (Sigma) 1:20, anti-Ubiquitin Ubi-1 ab7254 (abcam) 1:500, anti-DRP1 EPR19274 (abcam) 1:250, anti-TOM20 sc-17764 (Santa Cruz) 1:200. Secondary antibodies coupled to Alexa Fluor 488 (Invitrogen) or Cy3 (Jackson Immuno Research Laboratories, Inc.) were used at 1:300– 1:500. For Mito Tracker−Red CMXRos (Invitrogen) staining, cells were incubated during 45 min at 37°C with 400 nM Mito Tracker and washed three times with conditioned media prior to PFA fixation.

FISH for ribosomal RNA was performed using digoxigenin-labelled riboprobes as previously described (Thomas et al., 2005). Single-molecule FISH with Stellaris probes (Biosearch Technologies) was performed according to manufacturers’ instructions.

Labelling of translating polypeptides was performed by incubating the cells in complete medium with 10 nM puromycin (SIGMA) during 10 min, followed by fixation and detection with anti-puromycin (PMY-2A4, DSHB) and Alexa Fluor 488 secondary antibody. Fluorescence intensity in single cells was measured using ImageJ, (https://imagej.nih.gov/ij/) and average from 100 cells was calculated for each experimental point in duplicate coverslips.

Images were acquired with PASCAL-LSM, LSM510 Meta, or LSM 880 confocal microscopes (Carl Zeiss), using C-Apochromat 40×/1.2 W Corr or 63×/1.2 W Corr water immersion objectives for the PASCAL-LSM, an EC “Plan-Neofluor” 40×/1.30 NA oil or Plan-Apochromat 63×/1.4 NA oil objective lenses for the LSM510 Meta, and a Plan-Apochomat 63x/1.4 Oli DIC M27 objective for the LSM880. Pixel intensity was always lower that 250 (saturation at 255). Cell contours in Figures 1B and 1E were manually drawn using saturated immunofluorescence images as templates.

### Image analysis

To evaluate the presence of Smaug1-bodies or Smaug1-EYFP bodies in fixed cells, cells were manually classified as positive when more than three bodies of at least 0,25 µm in diameter were present. For smFISH confocal images, we used a previously described strategy (Luchelli et al., 2015) to evaluate the frequency of stochastic contacts of SDHB and UQCRC1 mRNAs with Smaug1 ECFP-bodies. Three cognate images of the Smaug1-body channel were generated by flipping and/or mirroring the original and colocalization was manually inspected. A total of 400 Smaug1-ECFP bodies from randomly-selected cells were analyzed and 3 to 7 ROIs covering almost all the cell cytosolic area were analyzed. Contacts with mitochondria were manually evaluated and the frequency of random events was evaluated in randomized images of Smaug1. At least five rectangular ROIs of 50 µm^2^ per cell were randomly selected, where mitochondria signal covered 30% to 50%.

### Confocal live-cell imaging

A Carl Zeiss LSM 880 inverted confocal microscope equipped with a stage-top heated platform maintained at 37°C and with a controlled CO_2_ flux chamber was used. U2OS cells were grown and transfected in 8 wells Nunc® Lab-Tek chamber slides. U2OS cells expressing hSmaug1-EYFP and Mito-DsRed were imaged 24 hs after transfection. Metformin was injected to slides without opening the CO_2_ chamber. Z-stack images were obtained for each cell every two minutes. For image analysis, acquired z-stacks were processed with a custom-made Python script to segment and quantify the intensity and morphological properties (Malik-Sheriff et al., 2018) of the Smaug1-EYFP bodies. The segmentation pipeline consisted in filtering the hSmaug1-EYFP channel with Laplacian of Gaussian filters and using manual thresholding to distinguish between pixels corresponding to Smaug1-EYFP bodies from background pixels. Smaug1-body pixels were later disregarded to distinguish between background and cytoplasm pixels. To analyze the effect of metformin, the number of Smaug1-EYFP bodies at each time point was normalized to the pre-treatment value for each cell. Afterwards, quantifications were binned every 4 minutes and averaged. For error estimation, error of the mean was used. For all the analysis, Python 3.7, numpy 1.15, pandas 0.23, scikit-image 0.14, and scipy 1.1 were used.

### Western blot

WB were performed by standard procedures using polyvinylidene fluoride membranes (Immobilon-P polyvinylidene difluoride, Millipore), ECL Prime (GE Healthcare) and analyzed using a LAS4000 Imager (GE Healthcare). Primary antibodies were used as follows: anti-SAMD4A HPA043061 (Sigma) 1:100; anti-SAMD4B HPA059385 (Sigma) 1:100, mouse anti-V5 (Invitrogen) 1:5,000, anti-GFP (Invitrogen) 1:2,000; anti-tubulin (DSHB) 1:10,000, anti-β-actin (Sigma-Aldrich) 1:10,000, anti-DRP1 EPR19274 (abcam) 1:1000, anti-OPA1 EPR11057(B) (abcam) 1:1000, anti-Mitofusin 2 6A8 (abcam) 1:1000, anti-TOM2O sc-17764 1:100. HRP-conjugated secondary antibodies were used 1:10,000 or 1:100,000. Signal intensity was assessed with the ImageJ software.

### Respiratory parameters and mitochondrial membrane potential

Oxygen flow in U2OS cells under ADP excess (state 3) was measured using a two-channel, high-resolution Oxygraph respirometer (Oroboros, Innsbruck, Austria), as described in (Doerrier et al., 2018). Briefly, cells were treated with the indicated siRNAs during 48hs and harvested at 90% confluence in trypsin-EDTA, washed once with phosphate buffered saline (PBS), resuspended in fresh growth media without antibiotics at a density of 1 × 10^6^ cells/ml, and analyzed in 2 ml Oxygraph chambers under continuous stirring at 750 r.p.m. at 37°C in 2 s intervals. For intact cell analysis, routine respiration was recorded and then 2.5μM oligomycin were added to inhibit the ATP synthase, which allows to measure leak respiration. The electron transfer system (ETS) capacity was evaluated by titration with carbonyl cyanide m-chlorophenyl hydrazone (CCCP) uncoupler in 0.5 μM steps until a maximum flow was reached. Respiration was inhibited by 0.5 μM rotenone and 2.5 μM antimycin A to determine residual oxygen consumption (ROX). For digitonin-permeabilized cells, a substrate-uncoupler-inhibitor-titration protocol termed SUIT-RP2 (Doerrier et al., 2018) was performed. Oxygen flow was measured before digitonin treatment in respiration medium MiR05 (Routine), and after successive addition of substrates and inhibitors as follows: 5 mM ADP, 0.1 mM malate, 0.2 mM octanoyl-carnitine, to record the electron-transferring flavoprotein complex from fatty acid b oxidation to coQ (F pathway); 2 mM malate, 5 mM pyruvate, 10 mM glutamate to record OXPHOS in the N-junction (Frostner et al., 2019) 10 mM succinate to record the OXPHOS capacity state FNS with convergent input of electrons via complexes I and II into the respiratory system, 10 mM glycerol-3-phosphate to activate the glycerophosphate dehydrogenase shuttle; stepwise titration with CCCP in 0.5 μM increments as needed to determine the ETS capacity state at maximum oxygen flow; 0.5 μM rotenone for complex I inhibition. All respiratory coupling states were corrected for ROX, which was obtained after the addition of 2.5 µM antimycin A. Finally, 5mM ascorbate plus 0.5mM N,N,N′,N′-tetramethyl-p-phenylendiamine dihydrochloride (TMPD) were used as substrates to assess complex IV activity. Mitochondrial membrane integrity was verified after addition of 10 μM cytochrome c, and changes were always lower than 10%. Data was analyzed using DatLab7.4.0.4 software (Oroboros, Austria). Oxygen flow is expressed as picomoles per second per million cells and normalized to the mitochondrial DNA content in each sample (pmol/ S*millcells*mtDNA). All experiments were performed using instrumental background correction and after calibration of the polarographic oxygen sensors. Statistical significance was calculated using Mann-Whitney rank-sum test using SPSS IBM for Windows statistical package, version 26 (SPSS Inc, Chicago, IL, USA)

Mitochondria depolarization was assessed with the MitoProbe 5’,6,6’-tetrachloro-1,1’,3,3’-tetraethylbenzimidazolylcarbocyanine iodide (JC-1). For cytometry assay, cells were treated with trypsin, resuspended in complete DMEM, washed twice with PBS and incubated with 2 μM JC-1 dissolved in pre-warmed PBS at 37 °C, 10% CO2 during 20 min and analyzed by flow cytometry.Exposure to 100 µM CCCP to induce complete depolarization was done during 5 min before incubation with JC-1. Flow cytometry analysis was performed using a FACS Aria (Becton Dickinson, Franklin Lakes, NJ, USA) and data mining was done using FlowJo (Tree Star, Inc., Ashland, OR, USA). Fluorescence intensity of J-aggregates (red) and JC-1 monomers (green) was measured in the mCherry and FITC channels. CCCP-treated cells were used to set gates. Geometric means normalized to siNT-treated cells were calculated. Confocal live-cell imaging was performed using a Carl Zeiss LSM 880 inverted confocal microscope as described above. Cells were seeded in Lab-tek chamber slides and incubated with JC-1 and CCCP as above. Images were obtained using a 20X objective and 488/ 543 nm lasers and fluorescence intensity was analyzed using ImageJ.

### Pull-down assays

Cells were grown in 100 mm-plates and transfected with a mix of 250 ng of VSBP-V5 tagged constructs and 250 ng of EYFP constructs. After 24 hours of expression, cells were harvested with 500 µl of NET buffer (150 mM NaCl, 50 mM Tris pH7.5, 1 mM EDTA and protease inhibitor cocktail from Sigma). Cells were kept on ice for 15 minutes and lysed by 20 times up and down pipetting. After a 15-min centrifugation to remove cell debris the supernatant (input, IN) was incubated for 1 hour at 4°C with Streptavidin-conjugated beads. After three washes in NET buffer supplement with 0.1% TX100 and one wash in NET buffer, pulled-down proteins were cracked in protein sample buffer and subjected to western blot for EYFP and/or V5.

### Tethering assays

Tethering assays were performed as previously described (Bhandari et al., 2014; Deng et al., 2015). In brief, HEK293T cells were transfected at 50% confluency with a mix of 250 ng of Smaug1-MS2-HA constructs, 200 ng of Firefly-6xMS2bs plasmid or control firefly and 50 ng of pCIneo-Renilla for 24 hs in 24-wells plates using Jet Prime reagent following the manufacturer’s instructions. Luciferase expression levels were analyzed using the Dual Luciferase kit (Promega) in a DTX880 Multimode detector (Beckman Coulter).

### RNA pull-down assays

Cells grown in 100 mm plates were transfected with 10 µg of V5-SBP tagged constructs and harvested 24 hs afterwards in 400 µl of RPD buffer (150 mM NaCl, 25 mM Tris pH7.4, 5 mM EDTA, 0.5 mM DTT, 1% NP40, 5% glycerol and protease inhibitor cocktail from Sigma). Cells were lysed using a Bioruptor (Diagenode) and after centrifugation, pull-down was performed with Streptavidin-conjugated beads. Total RNA from pull-down or input samples was isolated using TRIzol LS reagent (Invitrogen) following the manufacturer’s instructions. First-strand cDNA was synthesized from 2 µg of total RNA using random hexamers and MMLV reverse transcriptase (Promega). The cDNA was used as a template for quantitative PCR performed using Syber Green reagent (Applied Biosystems) and the Light Cycler 480 system (Roche). The amount of the indicated mRNAs relative to RpLp0 was determined using specific primers. Binding is expressed as the relative amount of mRNA in the pull-down material normalized to the relative amount in the input sample. The levels of the tagged constructs in the inputs and pulled-down material were measured by western blot using anti-V5 antibody. Sequences of primers were as follows: NDUFA10 Fw. AGTACTCAGATGCCTTGGAG and Rv, GCTCCAACACAACACCTTGTC; UQCRC1 Fw, CCTCTCAGCCCACTTGCA and Rv, CGGCTGCCAACATCAAT; SDHB Fw, GTGGCCCCATGGTATTGGAT and RV, CACAAGAGCCACAGATGCCT; PPIA Fw, TTCATCTGCACTGCCAAGAC and Rv TCGAGTTGTCCACAGTCAGC; RpLP0 Fw, GGGCAAGAACACCATGATGC and Rv CATTCCCCCGGATATGAGGC.

### Statistics

Each experimental point included duplicate or triplicate coverslips or wells. Cell numbers are indicated in each figure panel. Statistical significance was determined using Excel or Instat software (GraphPad Software, Inc.) in all figures unless indicated. P-values (*, P < 0.05; **, P <0.01; ***, P < 0.001; ns, not significant) relative to control treatments were obtained by one-way ANOVA and Tukey post-test. Error bars represent standard deviation from duplicate coverslips or from independent experiments, as indicated.

## Acknowledgements

We dedicate this work to Elisa Izaurralde (1959–2018). We thank Catia Igreja and Dipankar Bhandari, from Elisa Izaurralde’s laboratory (Max Planck Institute for Developmental Biology, Tübingen, Germany) for their generous advice on tethering and pull-down assays. We deeply thank Anabella Srebrow (IFIBYNE-UBA-CONICET, Argentina) for her generous support. We are grateful to Andrés Rossi (Instituto Leloir) for assistance in confocal imaging and to Marcelo Perez-Pepe for advice in puromycilation assays. This work was supported by the following grants : PICT 2013-3280 to GLB; PICT 2014-3658 to GLB and HG; PICT 2013-1301 to HG; PICT 2015-1302 to AJFA; PICT 2012-2493 to MGT, all them from ANPCyT (Argentina); PIP2011-205- (CONICET, Argentina) to MGT; an Alicia Moreau Chair from Paris Diderot University (France) to GLB and MINCYT-ECOS SUD A14S03 to GLB and AP; grant 1112 from the Fondation ARC pour la Recherche sur le Cancer to AP, and SAF2016-75004-R (MINECO, Spain) and PROMETEO/2018/055 (Generalitat Valenciana, Spain) to MC. MC participates in the COST Action CA15203 MITOEAGLE. AJFA, MGT, HG and GLB are career investigators from CONICET, GLB and HG are professors at the University of Buenos Aires, Argentina; MLP, MH, PELS, and AAC are or were recipients of PhD or Postdoctoral research fellowships from CONICET, JP was recipient of a CONICET PhD fellowship and a visiting doctoral fellow at the Carmo Fonseca Lab supported by H2020-Marie Sklodowska-Curie Research and Innovation Staff Exchanges [734825-LysoMod], AP is a professor at the University Paris Diderot.

## Author contributions

AJFA, MGT, MLP, HEG, MC, JP and GLB conceived and designed experiments and analyzed data. AJFA, MGT, MLP, MH, JP, MC, PELS, LB, JPP, AAC, MCF and GLB performed the experiments and/or analyzed data. All authors discussed the results; GLB conceived the study, supervised the project, and wrote the manuscript with the contribution of AJFA, MGT, HEG, MC and AP.

## Supplementary material

### Videos

**Video 1**. (Linked to Figure 1C). A representative Smaug1-EYFP transfected cell was recorded during 30 min as described in Materials and Methods.

**Video 2**. Live cell imaging of the control cell depicted in Figure 10D. Smaug1-EYFP (green) and Mito-dsRED (magenta) were co-transfected.

**Video 3**. Live cell imaging of the metformin-exposed cell depicted in Figure 10D. Smaug1-EYFP (green) and Mito-dsRED (magenta) were co-transfected.

**Supplemental Figure 1. Smaug1-bodies exclude ubiquitin and small ribosomal subunits**. Top, Smaug1-V5-transfected U2OS cells were immunostained for V5 and ubiquitin. Bottom, Smaug1-EYFP-transfected U2OS cells were submitted to FISH with a ribosomal 18S riboprobe as described in Material and Methods. Both markers are excluded from Smaug1-bodies.

**Supplemental Figure 2. Respiratory capacity of Smaug1+2-knockdown U2O2 cells. (A)** U2OS were treated with non-targeting (NT) siRNA or with siRNAs against both Smaug1 and Smaug2 (siS1+S2) and respiration in intact cells (1 × 10^6^ cells/mL) was examined in growth medium at 37 °C as described in Materials and Methods. Oxygen flow (pmol O_2_ × s^−1^ × 10^−6^ cells) and total oxygen concentration (nmol/mL) in the Oxygraph chamber are indicated as red and blue traces, respectively. CE: cellular substrate, P: Pyruvate, O: Oligomycin, U: Uncoupler (CCCP) R: Rotenone, A: Antimycin A. **(B)** After measuring routine oxygen consumption, the adenosine triphosphate (ATP)-synthase inhibitor oligomycin was added to evaluate proton LEAK. The uncoupler carbonyl cyanide m-chlorophenylhydrazone (CCCP) allowed the measurement of maximal oxygen consumption (Max) stimulating maximal respiration assuming all required substrates are present. Finally, rotenone (complex I inhibitor) and antimycin A (complex III inhibitor), which completely prevent oxygen consumption through the electron respiratory chain were added. Oxygen flow per cells was corrected for ROX at the indicated mitochondrial respiration state. Calculated mitochondrial (mt) flux control ratios show basal cellular routine respiration (R), leak respiration (L) and fraction of respiration (netR = R-L) used for ATP production normalized to ETS capacity.

**Supplemental Figure 3. Mitochondrial network disruption upon Smaug1 and Smaug2 knockdown**. U2OS cells were treated with the indicated siRNAs and live-stained with MitoTracker™ Red CMXRos. At least 200 cells per treatment were analyzed and the percentage of cells with elongated mitochondria is indicated. A representative experiment out of three is shown. Scale bars, 10 μm.

