## Supplemental Figure 1 for "Smaug membraneless organelles regulate mitochondrial function"

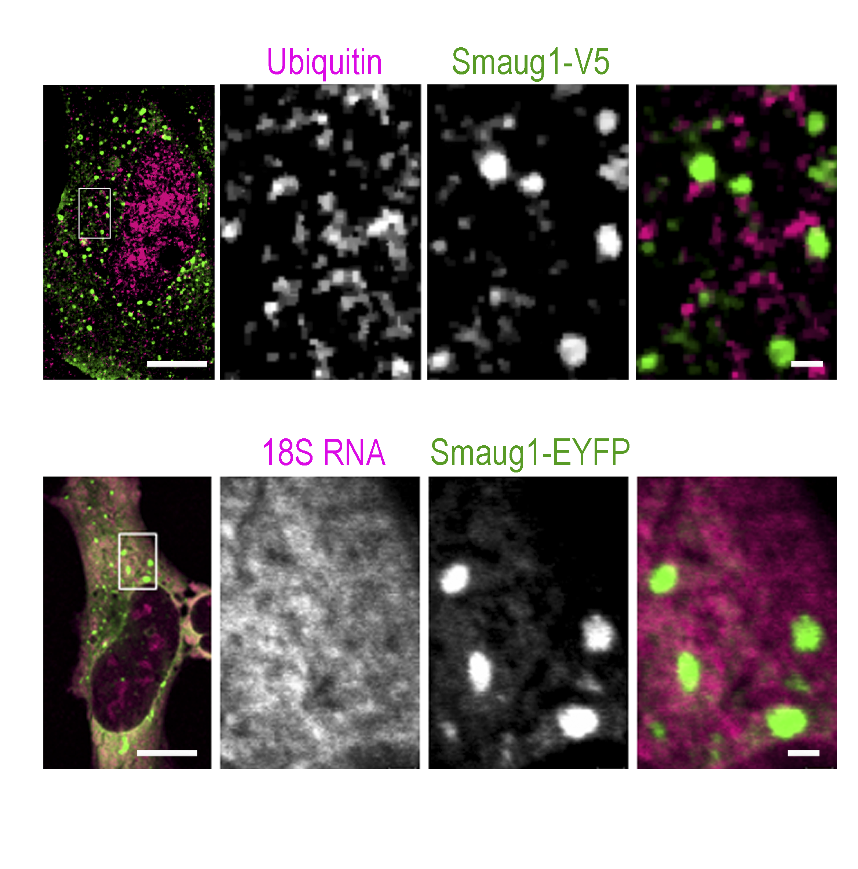

**Supplemental Figure 1. Smaug1-bodies exclude ubiquitin and small ribosomal subunits.**

Smaug1-V5-transfected U2OS cells were immunostained for V5 and ubiquitin. Smaug1-EYFP-transfected U2OS cells were submitted to FISH with a ribosomal 18S riboprobe as described in Material and Methods. Both markers are excluded from Smaug1-bodies
