## Supplemental Figure 2 for "Smaug membraneless organelles regulate mitochondrial function"

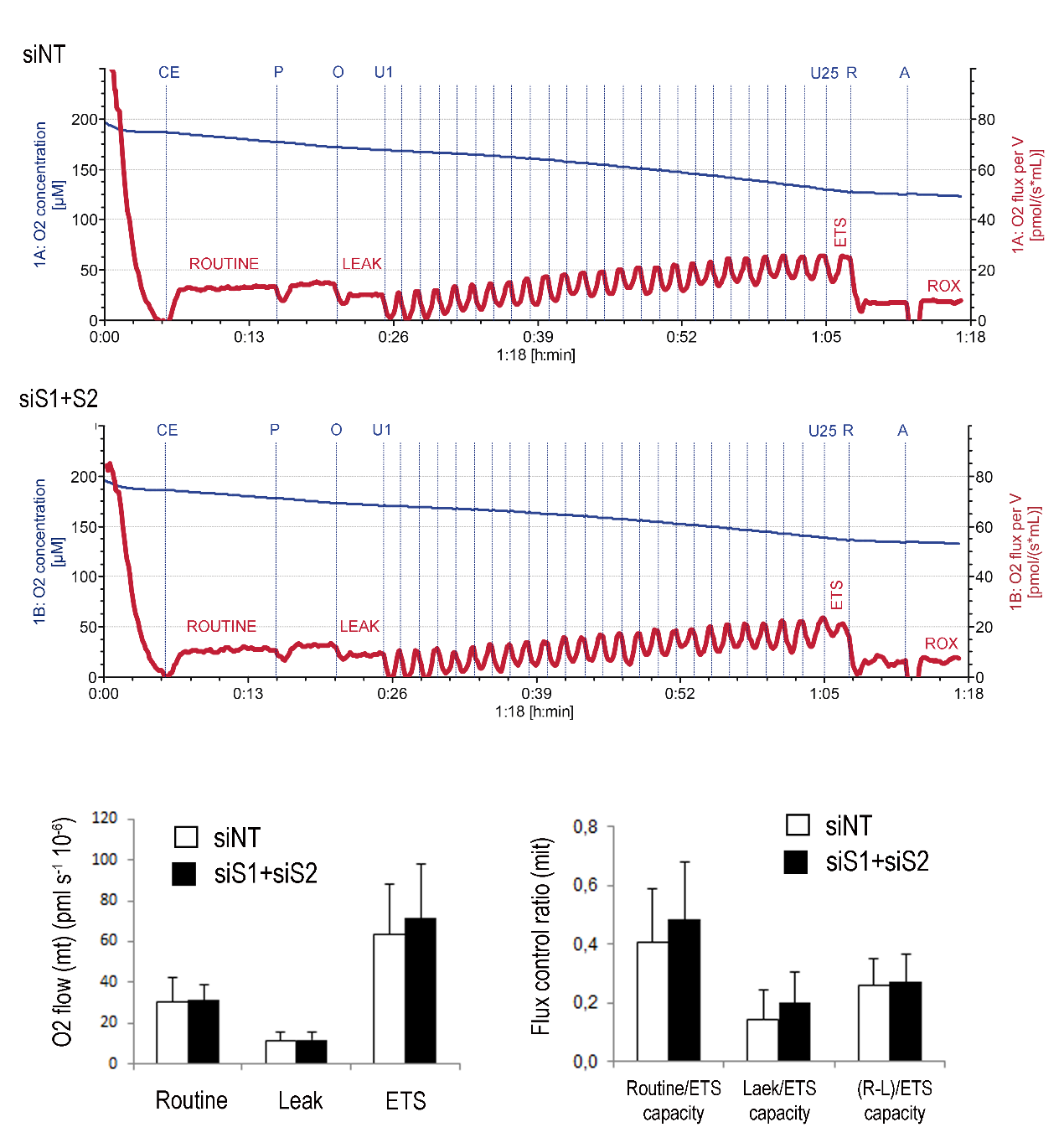

**Supplemental Figure 2. Respiratory capacity of Smaug1+2-knockdown U2O2 cells.**

**(A)** U2OS were treated with non-targeting (NT) siRNA or with siRNAs against both Smaug1 and Smaug2 (siS1+S2) and respiration in intact cells (1 × 10^6^ cells/mL) was examined in growth medium at 37 °C as described in Materials and Methods. Oxygen flow (pmol O_2_ × s^−1^ × 10^−6^ cells) and total oxygen concentration (nmol/mL) in the Oxygraph chamber are indicated as red and blue traces, respectively. CE: cellular substrate, P: Pyruvate, O: Oligomycin, U: Uncoupler (CCCP) R: Rotenone, A: Antimycin A. **(B)** After measuring routine oxygen consumption, the adenosine triphosphate (ATP)-synthase inhibitor oligomycin was added to evaluate proton LEAK. The uncoupler carbonyl cyanide m-chlorophenylhydrazone (CCCP) allowed the measurement of maximal oxygen consumption (Max) stimulating maximal respiration assuming all required substrates are present. Finally, rotenone (complex I inhibitor) and antimycin A (complex III inhibitor), which completely prevent oxygen consumption through the electron respiratory chain were added. Oxygen flow per cells was corrected for ROX at the indicated mitochondrial respiration state. Calculated mitochondrial (mt) flux control ratios show basal cellular routine respiration (R), leak respiration (L) and fraction of respiration (netR = R-L) used for ATP production normalized to ETS capacity.
