## Supplemental Figure 3 for "Smaug membraneless organelles regulate mitochondrial function"

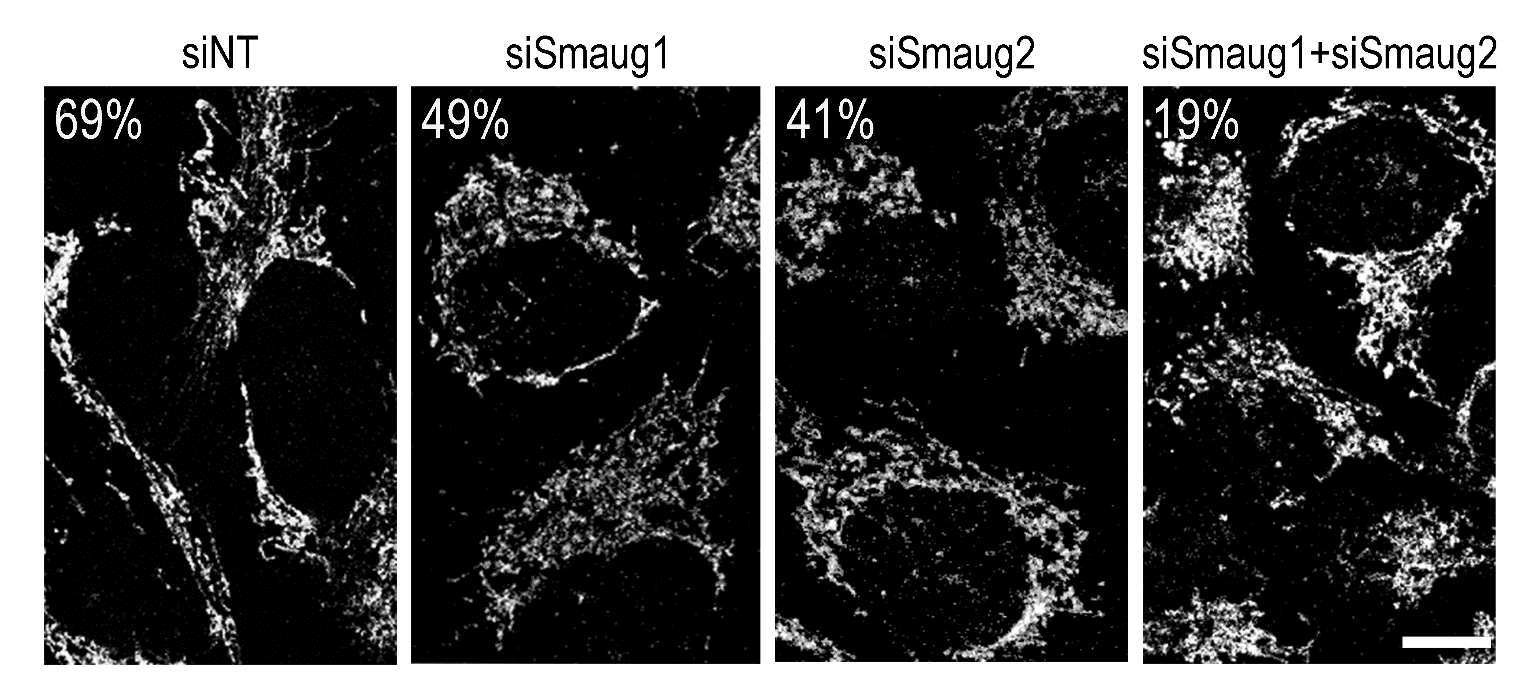

**Supplemental Figure 3. Mitochondrial network disruption upon Smaug1 and Smaug2 knockdown.** U2OS cells were treated with the indicated siRNAs and live-stained with MitoTracker™ Red CMXRos. At least 200 cells per treatment were analyzed and the percentage of cells with elongated mitochondria is indicated. A representative experiment out of three is shown. Scale bars, 10 μm.
